# Lymphatic networks enable diversification of B cell responses to vaccines

**DOI:** 10.64898/2026.09.16.752166

**Authors:** Jaime L Chao, Christopher D Thouvenel, Courtney E McDougal, Mark D Langowski, Neil P King, Marion Pepper, David J Rawlings, Michael Y Gerner

**Affiliations:** Department of Immunology, University of Washington School of Medicine, Seattle, WA, USA; Center for Immunity and Immunotherapies, Seattle Children’s Research Institute, Seattle, WA, USA; Department of Biochemistry, University of Washington, Seattle, WA, USA; Institute for Protein Design, University of Washington, Seattle, WA, USA

## Abstract

Vaccination generates heterogenous B cell responses that collectively provide protective immunity, yet how this diversity is established remains poorly understood. Here, we identify the draining lymphatic network as a spatial organizer of humoral immunity. Vaccine dispersal across interconnected draining lymph nodes (dLNs) establishes gradients of antigen and inflammation that create distinct B cell differentiation and selection environments. dLNs proximal to the vaccination site preferentially support antibody-secreting cells and clonally diverse germinal center (GC) responses. In contrast, distal dLNs impose more stringent selection, favoring clonally restricted GCs enriched for high-affinity B cells. Spatially distinct inflammatory environments further determine antibody isotype, linking LN position to antibody effector class. These spatial biases persist upon recall and can be reprogrammed through vaccine design, including nanoparticle properties and LN-directed targeting. Thus, lymphatic networks diversify B cell fate, repertoire and affinity selection, and antibody isotype, revealing a programmable axis for tuning humoral immunity through vaccine design.

## INTRODUCTION

B cell-mediated immunity is critical for protection against pathogens and depends on the coordinated generation of antibody-secreting cells (ASCs) that secrete large quantities of antibodies, GC B cells that evolve diverse and high affinity responses, and memory B cells (MBCs) that mediate rapid anamnestic recall responses. Generation of these functionally distinct populations from individual antigen-specific B cells is governed by the integration of multiple signals, whose magnitude, timing, and context collectively determine cell fate decisions.^1^ Antigen availability drives the magnitude of B cell expansion, while B cell receptor (BCR) signal strength and synergistic engagement of innate receptors, such as Toll-like receptors (TLRs), promotes early ASC differentiation.^2–8^ Within germinal centers, repeated interactions with cognate T follicular helper (Tfh) cells drive affinity maturation through iterative rounds of clonal selection, while additional signals govern GC exit and differentiation into long-lived ASCs and MBCs.^9,10^ In parallel, inflammatory cues, including cytokines produced by Tfh cells and other immune populations, direct class-switch recombination to generate functionally specialized antibody isotypes.^11–13^ Together, these diverse inputs establish tunable thresholds for activation, differentiation, and selection, producing humoral responses with distinct magnitudes, affinities, clonal diversity, and effector functions for comprehensive immunological coverage and host defense.

While current models largely consider these processes within a single lymphoid organ, the instructive signals that drive humoral immunity can extend beyond one draining site. Following immunization, antigens and adjuvants drain via lymphatic vasculature across multiple interconnected dLNs, generating spatial gradients with varying antigen availability and inflammation that decrease with increasing distance from the site of immunization. These gradients occur both within individual dLNs^14–16^ and across the lymphatic chain^17–19^ and are a conserved feature of lymphatic physiology.^20–22^ Previous studies demonstrated that these gradients establish spatially distinct microenvironments that program divergent innate and T cell responses,^14–17^ with proximal dLNs preferentially promoting effector differentiation and distal dLNs favoring central memory precursor development.^17^ Whether these same spatially organized cues similarly regulate B cell fate decisions and thereby shape humoral immunity remains unknown.

Here, using mouse models of vaccination together with B cell tetramers, BCR sequencing, and high-parameter confocal microscopy, we find generation of divergent B cell responses across the draining lymphatic network. Although independent antigen-specific B cell responses arise across multiple dLNs, proximal and distal sites differ in ASC support, repertoire and affinity selection, and antibody isotype composition, indicating that each dLN functions as an independent yet coordinated contributor to humoral immunity. We demonstrate these spatial biases are programmable through vaccine design, revealing the lymphatic network as a previously underappreciated spatial axis of immune regulation that can be harnessed for next-generation vaccine strategies.

## RESULTS

### The lymphatic network comprises multiple productive sites of B cell priming

We first quantified antigen dissemination following footpad immunization using soluble fluorescent antigen (OVA) and LPS, which allows for robust quantification by microscopy. Four hours after immunization, antigen was readily detected throughout the draining lymphatic network, with highest abundance in the proximal popliteal LN (popLN) and declining abundance across distal iliac and inguinal LNs (**Figures S1A and S1B**). This gradient was evident both across whole tissue sections and within the follicular dendritic cell (FDC) network (**Figure S1C**). We next utilized flow cytometry to quantify particulate antigen uptake by polyclonal B cells following immunization with alum-absorbed fluorescent OVA and LPS, a clinically relevant vaccine formulation.^23^ Consistent with the imaging-based observations, a larger proportion of polyclonal B cells acquired antigen in proximal compared to distal dLNs (**Figure S1D**), demonstrating that vaccination establishes graded antigen availability across interconnected dLNs.

We next assessed whether distal dLNs could mount productive humoral immune responses. Nine days following immunization with alum-adsorbed OVA plus LPS, prominent antigen-specific B cell responses were detected in both proximal and distal dLNs, composed of OVA-tetramer-binding CD138+ ASCs and GL7+ GC B cells (**Figures 1A-1C**). Confocal microscopy confirmed both proximal and distal dLNs established canonical GCs displaying stereotypical organization of Bcl6^hi^ dark zones and light zones containing PD-1^+^ Tfh cells and FDCs (**Figure 1D**), which were sustained for at least two weeks (**Figures S2A-S2C**). Together, these data indicate that productive humoral responses are initiated throughout the lymphatic chain. To determine whether responses within distal dLNs reflected local priming rather than lymphocyte migration from proximal dLNs, mice were treated with FTY720 on days 7 and 8 post-immunization, early timepoints of B cell accumulation. Blocking lymphocyte egress did not alter OVA-specific GC or ASC numbers (**Figure S2D**), demonstrating that distal dLNs generate independent antigen-specific B cell responses.

**Figure 1.**
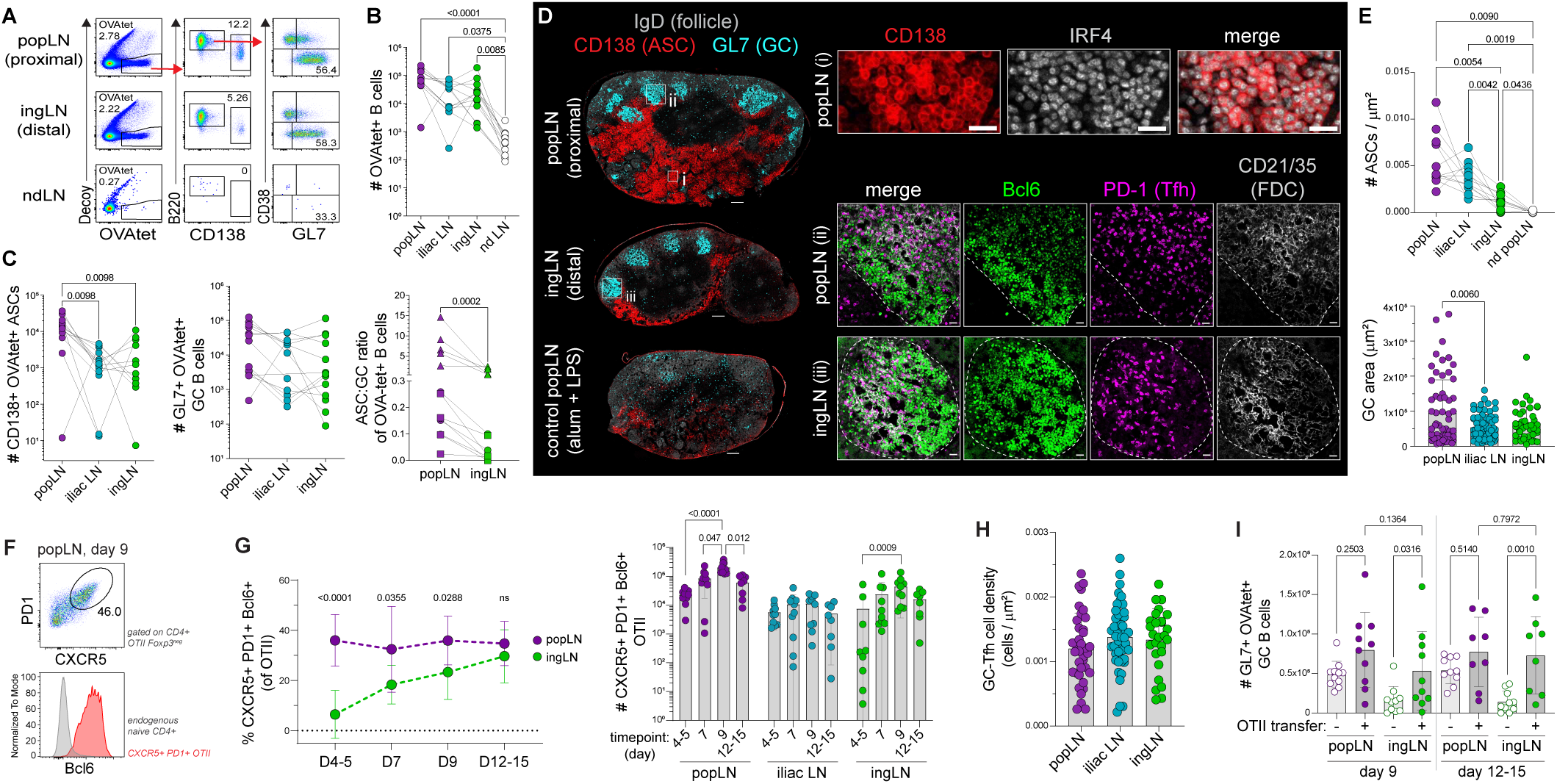
The lymphatic network comprises multiple productive sites of B cell priming. B6 mice were footpad immunized with alum-absorbed OVA and LPS; draining LN analyzed at day 9 unless otherwise indicated. A) Representative flow plots of CD19+ B cells in draining LNs. B-C) Flow quantification of OVA-tetramer+ B cells (B), and OVA-tetramer+ CD138+ ASCs, OVA-tetramer+ GL7+ GC B cells, and ASC:GC ratio among OVA-tetramer+ B cells (C, left to right). D) Representative confocal images of LNs depicting ASCs (inset i) and GCs (insets ii and iii). Scale bars = 200 μm (left), 20 μm (insets). E) Imaging quantification of IRF4+ CD138+ ASC density (top) and GC area (bottom). F) Representative flow plots of adoptively transferred CD45.1+ OT-II Tfh in popLN. G) Flow quantification of OT-II Tfh frequency (left) and number (right) at indicated timepoints. H) Imaging quantification of PD-1+ Tfh cell density within GC. I) Flow quantification of OVA-tetramer+ GC B cells at indicated timepoints, +/− OT-II adoptive transfer on day −1. Two-tailed one-way ANOVA with Dunn’s (B, C left and middle panels, G right panel, H, I), Tukey’s (E top panel), or Dunnett’s T3 (E bottom panel) multiple comparisons test, or Wilcoxon paired t-test (C right panel), or Mann-Whitney unpaired t-test (G left panel). Connected points indicate LNs from the same mouse (B, C, E), error bars represent s.d. Data are pooled from ≥2 experiments.

Despite these sites mounting productive humoral responses, proximal and distal dLNs exhibited quantitative and qualitative differences in cellular output. Proximal popLNs contained a trending increase in the number of tetramer-binding GC B cells compared to matched distal ingLNs (2.03-fold, **Figure 1C**), consistent with a subset of GCs displaying a larger cross-section area (**Figure 1E**). Notably, proximal popLNs exhibited a marked increase in the number of tetramer-binding ASCs (5.83-fold increase compared with distal ingLNs, **Figure 1C**), resulting in significantly elevated ASC:GC ratios among antigen-specific B cells (**Figure 1C**). Quantification of total CD138+ ASCs independent of tetramer binding confirmed this bias by both flow and microscopy, revealing higher ASC abundance and tissue density in proximal versus distal dLNs (**Figures 1D**, **1E, and S2E**) that persisted for at least two weeks after immunization (**Figures S2A-S2C**). Similar spatial bias of ASC responses in proximal popLNs was observed following immunization with alternative adjuvants (**Figures S2F and S2G**) and across a range of antigen doses (**Figure S2G**).

To determine whether these divergent ASC:GC responses were determined by position within the lymphatic network rather than LN identity, we altered the site of immunization to the tail base. In this model, the inguinal LN is proximal and the axillary LN is distal to the immunization site, while the popLN is non-draining.^19^ Under these conditions, antigen-specific B cell responses again occurred in both proximal inguinal and distal axillary dLNs, with preferential ASC differentiation occurring in proximal ingLN (**Figure S2H**). Together, these findings demonstrate that vaccination establishes multiple B cell priming sites across the draining lymphatic network. Rather than functioning as redundant units, individual sites exhibit distinct response biases and differ in their propensity to generate ASC responses.

### Delayed Tfh differentiation constrains germinal center responses in distal lymph nodes

We next investigated whether spatial gradients across the lymphatic network also regulate Tfh differentiation, a key determinant of GC responses. Endogenous CXCR5+ PD-1^HI^ Tfh phenotype cells were significantly reduced in distal ingLNs as early as day 5 after immunization, before GC formation (**Figure S2I**), suggesting divergent capacities of distinct sites to support early Tfh differentiation. Indeed, at 24 hours post-immunization, proximal dLNs contained elevated IL-6 concentration whereas distal dLNs exhibited increased IL-2 (**Figure S2J**), cytokine profiles associated with opposing ability to drive Tfh programming.^24–28^ To determine whether these differences persisted in T cells with uniform TCR affinity,^29^ we next examined monoclonal OVA-specific TCR transgenic CD4+ (OT-II) T cells following adoptive transfer. Four days post-immunization, robust OT-II Tfh cell formation occurred in proximal popLNs yet was significantly reduced in distal ingLNs (**Figures 1F and 1G**), mirroring observations of the endogenous polyclonal population. Notably, these differences did not persist, with the frequency and total number of OT-II Tfh cells progressively increasing in distal ingLN over time to match those in proximal popLNs (**Figure 1G**). Similarly, the density of endogenous polyclonal GC-Tfh cells (PD1+ CD4+ T cells in GCs) by day 9 was equivalent across proximal and distal sites (**Figure 1H**). Together, these findings reveal distinct kinetics of Tfh differentiation across the lymphatic network, with early timepoints favoring early Tfh differentiation in proximal dLNs and eventual convergence of GC-Tfh cell density over time.

We hypothesized that delayed Tfh differentiation reflects limited T cell help, resulting in restricted B cell responses in distal dLNs. To test this, we provided additional antigen-specific CD4+ T cells through OT-II adoptive transfer prior to immunization. This selectively enhanced the number of antigen-specific GC B cells in distal dLNs (**Figure 1I**), resulting in quantitatively similar GC outputs across the lymphatic network. Total ASC numbers were not significantly impacted at day 9, indicating that early ASC differentiation was largely unaffected. Instead, provision of additional cognate CD4+ T cells via OT-II T cell adoptive transfer resulted in increased ASC responses in distal ingLNs at day 15, a timepoint suggestive of GC-derived ASC output (**Figure S2K**). These findings indicate that delayed Tfh differentiation constrains early GC responses in distal dLNs.

### Distal draining lymph nodes impose stringent selection thresholds that shape germinal center clonal architecture

Because antigen availability and early Tfh differentiation both regulate affinity-dependent B cell recruitment to and selection within GCs,^1,9,10,30–32^ we next asked whether the spatial differences in these signals across the lymphatic network establish differential selection landscapes during early GC formation. To test this, we first assessed the avidity of B cell responses using the 4-hydroxy-3-nitrophenyl acetyl (NP) hapten system, in which probes of differing hapten valencies distinguish high-versus low-avidity GC B cells. Nine days after immunization, proximal and distal dLNs contained comparable numbers of high-avidity NP(8)-binding GC B cells (**Figures 2A and 2B**). In contrast, proximal popLNs recruited substantially more low-avidity NP(28)-binding GC B cells, resulting in a relative enrichment of high-avidity binders in distal ingLNs compared with matched proximal popLN (**Figures 2A and 2B**). Enrichment for high-avidity GC B cells at distal dLNs similarly occurred when an alternative lymphatic chain was targeted via tail base immunization (**Figure S3A**). Although the relative avidity increased at all sites between days 9 and 15, demonstrating affinity maturation occurs across the lymphatic network, the increase was greater in proximal dLNs (4.8-fold in proximal popLN versus 2.8-fold in distal iliac and ingLNs; **Figures 2B**, **2C and S3B**). These findings indicate that individual dLNs initiate GC responses under distinct selection thresholds, where proximal dLNs encompass a broader range of B cell avidities while distal dLNs are enriched for high-avidity GC B cells.

**Figure 2.**
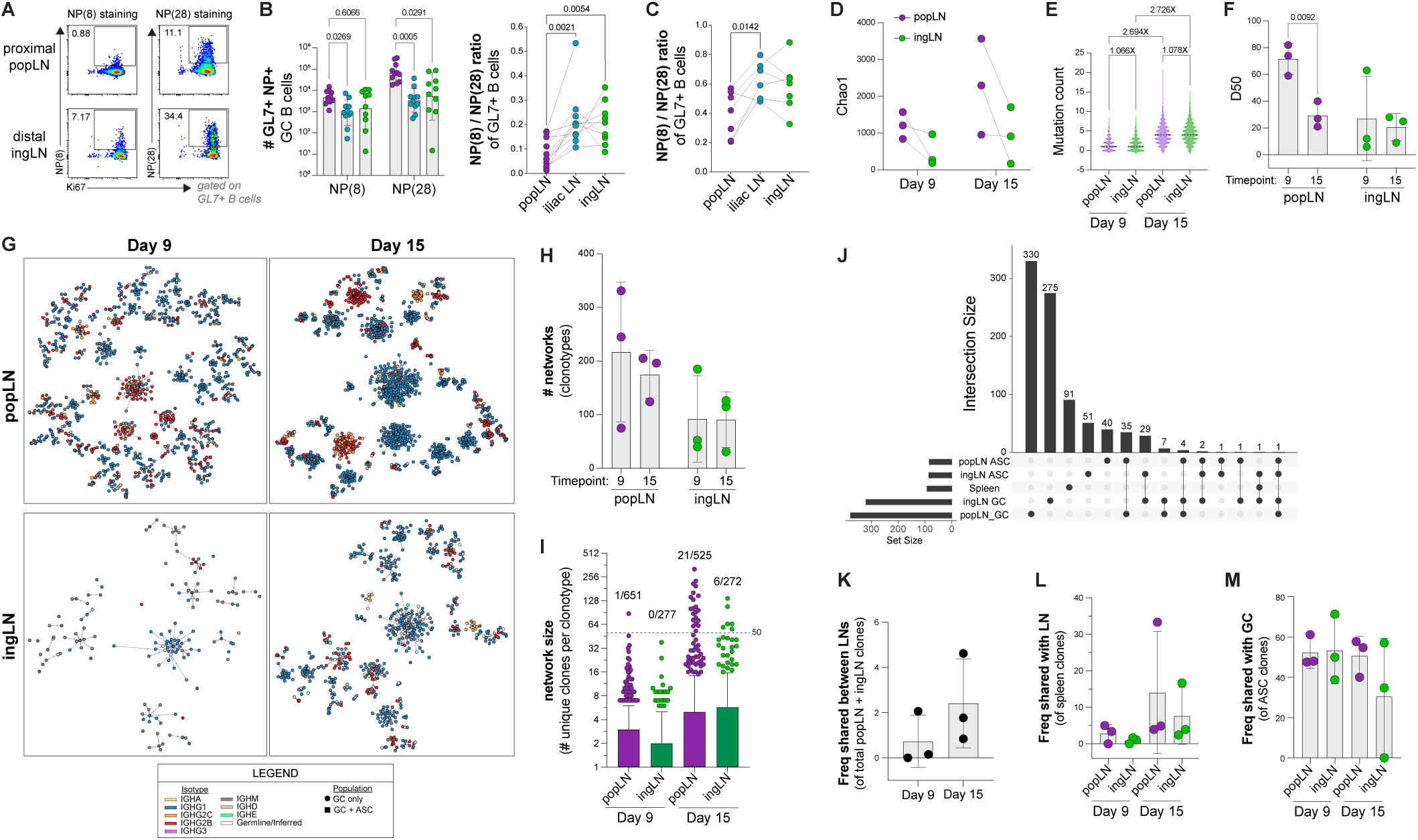
Proximal and distal draining LNs are defined by distinct repertoires. B6 mice were footpad immunized with alum-absorbed NP-OVA (A-C) or OVA (D-M) and LPS. A) Representative flow plots of GC B cells stained with NP(8) or NP(28). B-C) Quantification of total NP(8)-and NP(28)-binding GC B cells and NP(8)+/NP(28)+ GC B cell ratio at day 9 (B) and day 15 (C). D-M) BCR repertoire analysis of sorted OVA-tetramer+ populations at days 9 and 15 post-immunization. D) Chao1 estimator of repertoire diversity. E) Total mutation count per sequence. F) D50: minimum number of clones comprising 50% of total repertoire. G) Clonal network diagrams (each represents an individual LN). Each dot represents a unique sequence, connected to related sequences within same clonal family (clonotype network) based on nearest neighbor. Branch length proportional to mutation count. H) Number of clonotype networks per LN. I) Number of unique sequences per network. Proportions denote networks with ≥50 unique sequences (dotted line) over combined total networks for each LN at designated timepoint. J) Representative UpSet plot displaying repertoires of indicated populations from a single day 9 mouse. Horizontal bars = number of clones (set size) per population; vertical bars = intersection set size (black dots). K-M) Frequency of clones shared between popLN and ingLN (K), splenic clones also detected in the indicated LN (L), or ASC clones also detected as a GC clone within the same LN (M). Two-tailed one-way ANOVA with Dunn’s (B, left panel) or Tukey’s multiple comparisons test (B, right panel, C), or Welch’s unpaired t-test (F). Connected points indicate LNs from the same mouse (B-D), error bars represent s.d. (B, F, H, K-M), box-and-whisker plots represent 10-90 percentile (I). Data are pooled from 3 experiments (B, C), or representative of three independent mice per group (D-M).

These differences in GC population avidity suggested that proximal and distal dLNs may establish distinct clonal architectures. We therefore performed IgH BCR sequencing on sorted OVA-specific tetramer-binding GC B cells and ASCs isolated from matched proximal and distal dLNs, as well as all tetramer-binding cells from the spleen, at 9 and 15 days following OVA immunization (three mice per time point; 12,031 unique sequences analyzed; **Data Table S1**). Analysis of clonal abundance (Chao1) revealed that the GC population in proximal popLNs exhibited greater clonal diversity compared with matched distal nodes (**Figure 2D**), although this did not reach statistical significance, consistent with broader initial clonal recruitment in proximal dLN. Despite reduced clonal diversity, GC B cells within distal dLNs accumulated somatic mutations at comparable rates to those in proximal dLNs (**Figures 2E and S3C**), again indicating that both sites support productive affinity maturation.

We next asked whether the distinct initial clonal landscape affected the competitive structure of GC responses by examining repertoire evenness using D50 analysis (**Figure 2F**), which measures the number of dominant clones required to account for 50% of the repertoire. Proximal popLNs exhibited higher D50 values on day 9, consistent with broad and even participation of individual clones in the GC response. By day 15, however, D50 values declined in proximal popLNs despite maintaining a relatively high clonal abundance (**Figures 2D and 2F**), suggesting the emergence of dominant clones. In contrast, D50 values in distal ingLN remained comparability low at both timepoints, indicating initial recruitment of fewer clones that maintained a relatively stable distribution during GC evolution.

Clonal connectivity analysis confirmed these findings by visualizing the size of individual clonotypes within each LN repertoire (**Figure 2G**).^33^ Although the total number of clonotypes remained stable over time (**Figure 2H**), in line with stable estimated clonal abundance (**Figure 2D**), very large clonal networks (>50 unique sequences) preferentially emerged within proximal popLNs by day 15 (**Figure 2I**), indicative of unequal expansion of a few select clonotypes. In contrast, distal ingLNs maintained relatively stable clonal distributions over time (**Figures 2F-I**), reflecting more even participation of the smaller number of clones initially recruited into the GC response. Together, these findings indicate that proximal and distal dLNs support distinct modes of GC evolution. Proximal dLNs recruit broad repertoires that are progressively refined, suggestive of increasing inter-clonal competition, whereas distal dLNs establish smaller repertoires early that remain comparatively stable over time.

We next determined the clonal overlap of responses between matched proximal and distal dLNs within individual animals (**Figures 2J and S3D; Data Table S1**). Shared clones between both sites were rare, ranging from 0-4.62% of the combined repertoire (**Figure 2K**). This limited overlap indicates that individual dLNs generate largely independent B cell responses, consistent with our finding that blocking lymphocyte egress with FTY720 does not alter responses (**Figure S2D**). Notably, splenic clones could be detected in both proximal and distal dLNs (**Figure 2L**), demonstrating that both sites contribute to the systemic pool of antigen-reactive B cells.

Additionally, approximately 50% of ASC clones were also detected within matched GC repertoires at both day 9 and 15 timepoints (**Figure 2M**), demonstrating that a substantial fraction of ASCs originated from local GC responses. This relationship was independently validated on a cellular level using tamoxifen-inducible S1pr2-CreERT2-tdTomato GC fate-mapping mice.^34^ After three days of daily tamoxifen administration, up to 30% of ASCs in dLNs were tdTomato+ by day 9 (**Figure S3E**).

Collectively, these findings demonstrate that lymphatic gradients generate distinct GC selection environments across the draining network. Distal dLNs establish early GC repertoires enriched for high-avidity clones, whereas proximal dLNs recruit broader repertoires that include lower-avidity B cells and undergo oligoclonal expansion during GC evolution. Despite these distinct clonal trajectories, both sites support productive affinity maturation and contribute to systemic humoral immunity.

### Proximal draining lymph nodes establish expanded medullary niches that retain antibody-secreting cells

Since proximal popLNs were consistently enriched for ASCs following immunization (**Figures 1C**, **1E, S2B, S2C, and S2E-H**), we next asked whether these sites establish specialized microenvironments that support ASC accumulation. Indeed, early inflammatory cues differed across the lymphatic chain, with proximal popLNs exhibiting elevated levels of TNFα, IL-6, and IL-10 (**Figure S2J and S4A**), cytokines implicated in BLIMP1 induction as well as ASC differentiation, proliferation, and survival.^35–38^

Consistent with previous reports, the majority of ASCs localized to the LN medulla (**Figures 3A and 3B**).^35,39–41^ Notably, this medullary compartment underwent a transient yet substantial antigen-driven expansion selectively in proximal popLNs, occupying a significantly greater proportion of the total LN area by day 9 post-immunization (**Figures 3C and S4B**). ASC density correlated more strongly with medullary area than with total LN size (**Figure S4C**), identifying medullary expansion as a major determinant of local ASC accumulation.

**Figure 3.**
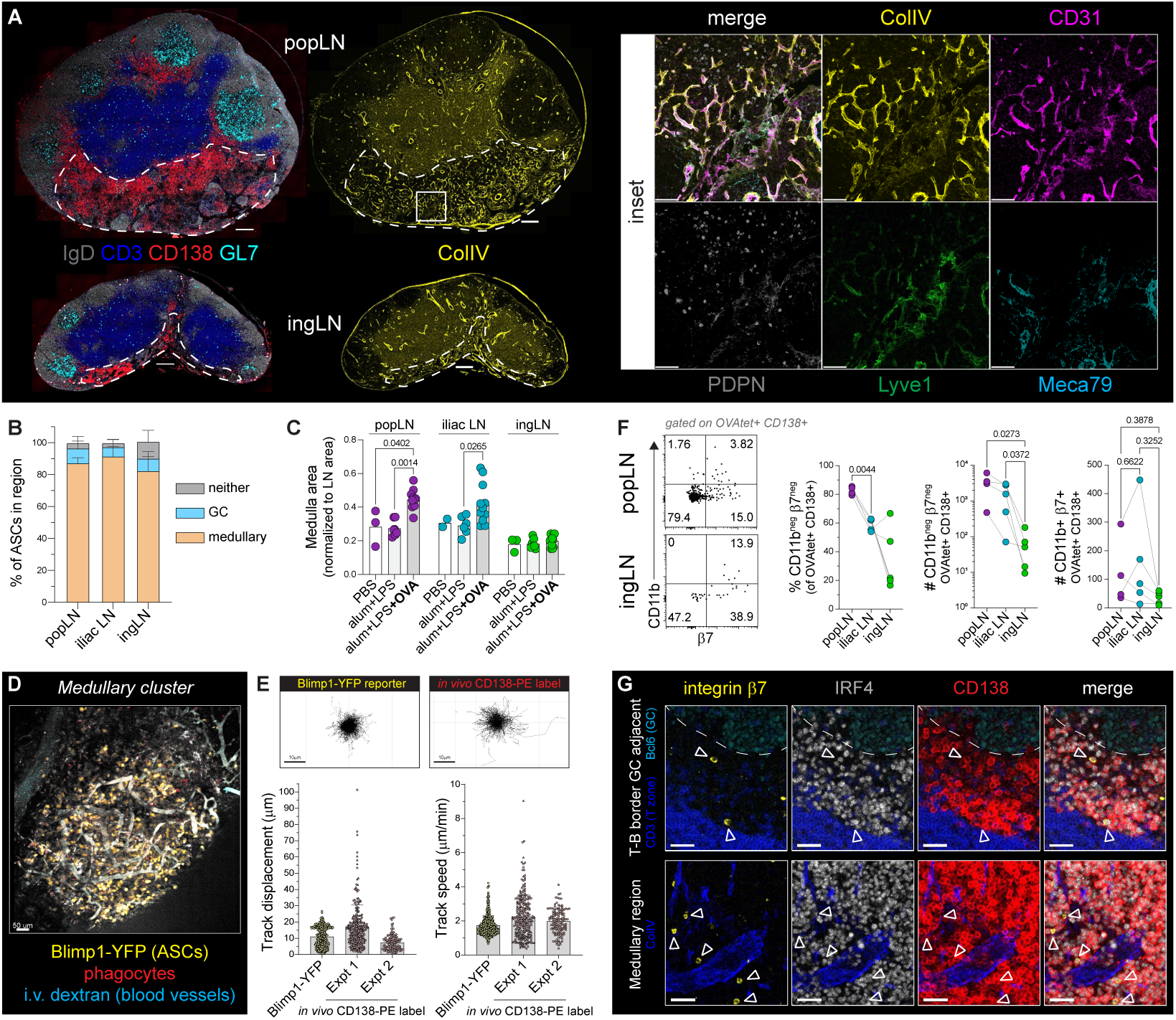
Proximal draining LNs establish expanded medullary niches that retain ASCs. B6 mice were footpad immunized with alum-absorbed OVA and LPS; draining LN analyzed at day 9. A) Representative images of draining LNs; outline denotes medulla. Scale bar = 200 μm (left) or 50 μm (insets). B) Imaging quantification of IRF4+ CD138+ ASC frequencies within the medulla, in or adjacent to GCs, or neither. C) Medulla area, normalized to total LN area. D) Representative intravital two-photon still-frame of a popLN from immunized Blimp1-YFP mouse. E) Intravital two-photon imaging quantification of ASCs in immunized Blimp1-YFP or *in vivo* CD138-PE-labelled B6 mice. (Top) Tracks of individual YFP+ ASCs or CD138-PE-labelled ASCs, centered at starting point. (Bottom) ASC track displacement and average speed. Symbols represent individual cells within an experiment. F) Flow quantification of OVA-tetramer+ CD138+ ASCs in draining LNs. Connected points indicate LNs from the same mouse. G) Representative confocal images of popLN depicting β7+ IRF4+ CD138+ ASCs (white arrowheads) at the T-B border adjacent to a GC (top) and within the medulla (bottom). Two-tailed one-way ANOVA with Dunn’s (C) or Tukey’s (F) multiple comparisons test. Error bars represent s.d. Data pooled from ≥2 experiments, or representative of ≥2 independent experiments.

Expansion of the medulla was associated with the development of extensive collagen IV-positive stromal networks (**Figure 3A**).^41^ Phenotypic analysis demonstrated that these structures consisted of CD31⁺ endothelial cells lacking expression of lymphatic endothelial markers PDPN and Lyve1, as well as the high endothelial venule marker Meca79. This phenotype is distinct from previously reported medullary reticular cells^40^ and instead is consistent with expanded capillary networks that may provide a specialized niche for ASC accumulation during the immune response.

To better understand the behavior of ASCs within these remodeled niches, we performed intravital two-photon microscopy. ASCs were visualized using either a Blimp1-YFP reporter or *in vivo* labeling with fluorescent anti-CD138 antibody. By day 9, ASCs formed dense clusters embedded within capillary-rich networks exhibiting active blood flow (**Figure 3D**; **Videos 1 and 2**), closely matching the structures observed by confocal microscopy (**Figure 3A**). Consistent with previous reports,^39,42^ ASCs exhibited minimal displacement and low motility (**Figure 3E**), indicating that they become largely sessile following localization within the medullary niche. Indeed, treatment with FTY720 to inhibit lymphocyte egress did not alter either total or antigen-specific ASC numbers within dLNs (**Figure S2D**), consistent with their sessile behavior and the minimal clonal overlap observed between proximal and distal dLNs (**Figure 2L**). Furthermore, the majority of antigen-specific ASCs in proximal dLNs lacked expression of CD11b and integrin β7 (**Figure 3F**), molecules required for LN egress and bone marrow homing.^43^ Together, these data indicate that most ASCs generated in proximal dLNs remain locally positioned within the expanded medullary niche rather than rapidly emigrating to distal sites.

Nevertheless, a subset of antigen-specific ASCs expressed both CD11b and β7, consistent with acquisition of bone marrow homing potential.^43^ These β7^+^ ASCs were broadly distributed throughout the LN medulla and were also detected at the T-B border adjacent to GCs (**Figure 3G**), suggesting that acquisition of BM homing potential can occur early after differentiation and GC exit. Consistent with this interpretation, a subset of ASCs exhibited substantially greater motility during two-photon imaging (**Figure 3E**), potentially representing these migratory CD11b^+^β7^+^ ASCs. Notably, despite the marked enrichment of total ASCs in proximal popLNs, the absolute number of antigen-specific CD11b^+^β7^+^ ASCs was comparable between proximal and distal dLNs (**Figure 3F**), suggesting that all activated dLNs within the lymphatic network can similarly contribute to the generation of ASCs with BM homing potential.

Collectively, these findings indicate that immunization establishes divergent medullary environments across the lymphatic network. Proximal dLNs undergo transient medullary remodeling associated with local accumulation and retention of largely sessile ASCs, whereas distal dLNs lack such expanded medullary architecture but still generate comparable numbers of ASCs with BM homing potential despite marked differences in total ASC output.

### Inflammation gradients regulate divergent antibody isotype profiles across proximal and distal draining lymph nodes

Inflammatory cytokines are well-established regulators of antibody class switching.^11–13^ Because proximal and distal dLNs experience distinct inflammatory environments following immunization (**Figures S2J and S4A)**,^17^ we hypothesized that local cytokine cues would differentially shape the antibody class switching events. Indeed, nine days after immunization, antigen-specific GC B cells exhibited distinct isotype profiles across the lymphatic network. IgG2b+ GC B cells were largely restricted to proximal popLNs, both by frequency and total number, whereas distal ingLNs contained higher frequencies of IgG1+ and IgM+ GC B cells (**Figures 4A**, **4B, and S5A**). Notably, the absolute numbers of antigen-specific IgG1+ and IgM+ GC B cells were comparable between proximal and distal dLNs (**Figures 4B and S5A**), indicating that their apparent enrichment in distal dLNs primarily reflected the selective accumulation of IgG2b+ and other isotype-switched populations within proximal dLNs.

**Figure 4.**
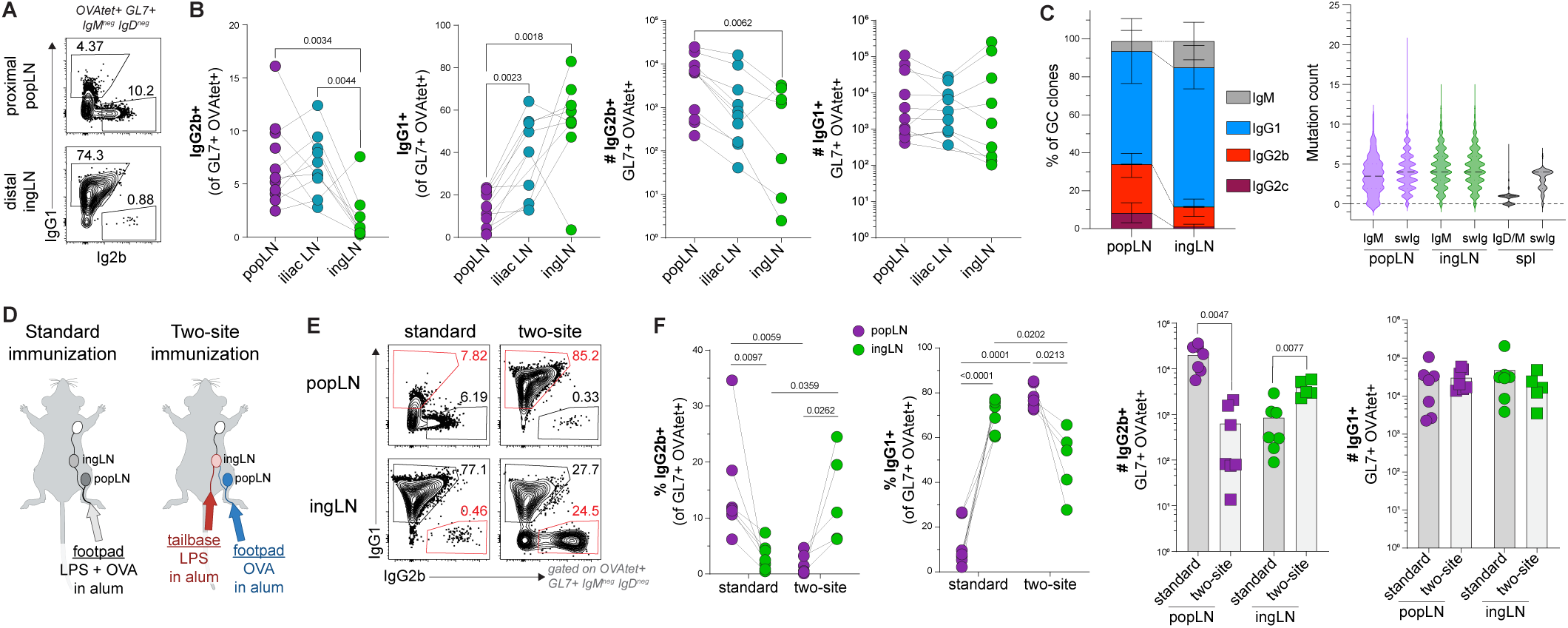
Inflammation gradients regulate divergent isotype profiles across proximal and distal draining LNs. A-C) B6 mice were footpad immunized with alum-absorbed OVA and LPS. A) Representative flow plots of OVA-tetramer+ GL7+ IgM^neg^ IgD^neg^ switched GC B cells, 9 days post-immunization. B) Day 9 flow quantification of IgG2b+ and IgG1+ OVA-tetramer+ GC B cells. C) Day 15 BCR repertoire quantification of C-gene usage within total OVA-tetramer+ GC clones (left); mutation count by indicated isotype in OVA-tetramer+ GC clones from draining LNs or total OVA-tetramer+ clones from spleen (right). D) Experimental setup for E-F. Mice were immunized in the footpad with alum-absorbed OVA and LPS and injected at the tail base with PBS (standard immunization), or immunized at both sites: in the footpad with alum-absorbed OVA and the tail base with alum-absorbed LPS (two-site immunization). E) Representative flow plots of OVA-tetramer+ GL7+ IgM^neg^ IgD^neg^ switched GC B cells 9 days post standard versus two-site immunization. F) Day 9 flow quantification of IgG2b+ and IgG1+ OVA-tetramer+ GC B cells after standard versus two-site immunization. Two-tailed one-way ANOVA with Tukey’s multiple comparisons test (B), or paired t-test (F left panels) or Welch’s unpaired t-test (F right panels). Connected points indicate LNs from the same mouse. Data are pooled from ≥2 experiments (B, F), or pooled from three independent mice per group (C).

These results were consistent with robust detection of IL-4, the primary driver of IgG1 switching, in both proximal and distal dLNs by day 7 (**Figure S4A**). Accordingly, IgG1+ responses were abolished in IL-4-deficient mice,^13,44^ whereas IgG2b+ responses in proximal dLNs remained intact (**Figures S5B and S5C**), indicating that the spatial distribution of antibody isotypes along the lymphatic network reflects underlying differences in cytokine requirements and availability. This spatial skewing of isotype profiles was maintained at later timepoints (**Figure S5D**), and was also observed following hapten-based immunization, occurring similarly in both high-and low-avidity populations (**Figure S5E**). Furthermore, the preferential bias toward IgG2b class-switching in proximal popLNs occurred with OVA + Sigma Adjuvant System (SAS) immunization (**Figure S5F**), suggesting that this spatially encoded heterogeneity of isotype class-switching is a generalizable principle of subunit vaccination.

Of note, many IgG1+ GC B cells in proximal popLNs displayed reduced surface IgG1 expression compared to those in distal ingLNs (**Figure 4A**). While intracellular Ig staining improved detection of IgG1+ cells (**Figures S5G and S5H**), a population of isotype switched antigen-specific GC B cells in proximal popLNs remained undefined, suggesting a broader isotype repertoire. To more comprehensively define antibody isotype usage, we analyzed IgH constant region representation within our BCR-sequencing dataset. Consistent with the flow cytometric analysis, proximal popLNs contained increased representation of IgG2b+ clones (**Figures 2H and 4C**). We also observed increased IgG2c representation, an isotype driven by IFNγ, in proximal popLNs, which was absent in distal ingLNs (**Figure 4C**). Conversely, distal ingLNs were enriched for IgM+ and IgG1+ clones (**Figure 4C**), mirroring flow-based analyses above. Notably, IgM+ GC clones accumulated mutations at levels comparable to the class-switched populations and above circulating naive IgD/M+ B cells isolated from the spleen (**Figure 4C**), suggesting that distal dLNs support affinity-matured IgM+ memory B cells. Together, these data indicate that distinct inflammatory environments across the lymphatic network shape local Ig isotype repertoires, generating spatially distributed and qualitatively diverse humoral outputs.

To directly test whether the local inflammatory landscape determines isotype profiles, we developed a two-site split immunization strategy that selectively redistributed inflammatory signals while preserving the original pattern of antigen drainage (**Figure 4D**). Mice were immunized in the footpad with alum-absorbed OVA plus alum to initiate responses in proximal popLN and distal ingLN. To manipulate the inflammatory gradient, LPS was either co-administered in the footpad (standard immunization) or delivered with alum at the tail base (two-site immunization), thereby selectively targeting LPS-induced inflammatory signals to the distal ingLN with no drainage to the proximal popLN.

Redirecting LPS-driven inflammation to the distal ingLN completely reversed the spatial distribution of IgG2b⁺ class switching. Consistent with earlier data, antigen-specific IgG2b⁺ GC B cells were enriched in the proximal popLN upon standard one-site immunization. However, upon two-site split immunization, IgG2b+ cells preferentially accumulated in the distal ingLN, increasing in both frequency and total number (**Figures 4E and 4F**). Conversely, antigen-specific IgG1⁺ GC B cells became enriched in the proximal popLN by frequency, whereas their total numbers remained largely unchanged (**Figures 4E and 4F**). Together, these findings demonstrate that local inflammatory environments along the lymphatic network dictate isotype composition and can be spatially reprogrammed by targeted delivery of inflammatory cues.

Two-site immunization also substantially reduced proximal ASC responses. In addition to a ∼3-fold reduction in total antigen-specific GC B cells, likely reflecting loss of IgG2b+ cells, total ASCs decreased by ∼26-fold, resulting in a reduced ASC:GC ratio in proximal dLNs compared to standard immunization (**Figure S5I**). These data suggested that the enhanced proximal ASC response following standard immunization is driven in part by local synergy between BCR and TLR signaling, a process known to promote GC-independent extrafollicular responses,^7,8,45^ and is consistent with an ASC response composed of both GC-independent and GC-derived populations (**Figures 2M and S3E**).

### Vaccine design can tune B cell responses at distal draining lymph nodes

Having established that lymphatic gradients regulate multiple aspects of humoral immunity, we next asked whether these spatial programs could be deliberately manipulated through vaccine design. Because transport of vaccine materials through the lymphatic vasculature is influenced by both physical delivery parameters and immunogen properties, we tested whether altering these features could selectively modulate B cell responses across the draining lymphatic network.

Bolus immunization generates local hydrostatic and osmotic forces that influence lymphatic fluid uptake and transport.^46^ We therefore hypothesized that increasing injection volume would enhance the delivery of vaccine components to distal dLNs. To test this, mice were immunized with either low-volume (10 μL) or high-volume (30 μL) injections while keeping antigen and LPS doses constant. Antigen-specific GC B cell responses in proximal popLNs were not impacted by injection volume 9 days post-immunization, indicating efficient delivery to the primary dLN under both conditions. In contrast, high-volume immunization significantly enhanced GC responses within distal ingLNs (**Figures 5A and 5B**), with marginal impact on ASC responses (**Figure S6A**), demonstrating that engagement of distal dLNs can be tuned by simple adjustments to injection volume without altering antigen or inflammatory agonist dosing.

**Figure 5.**
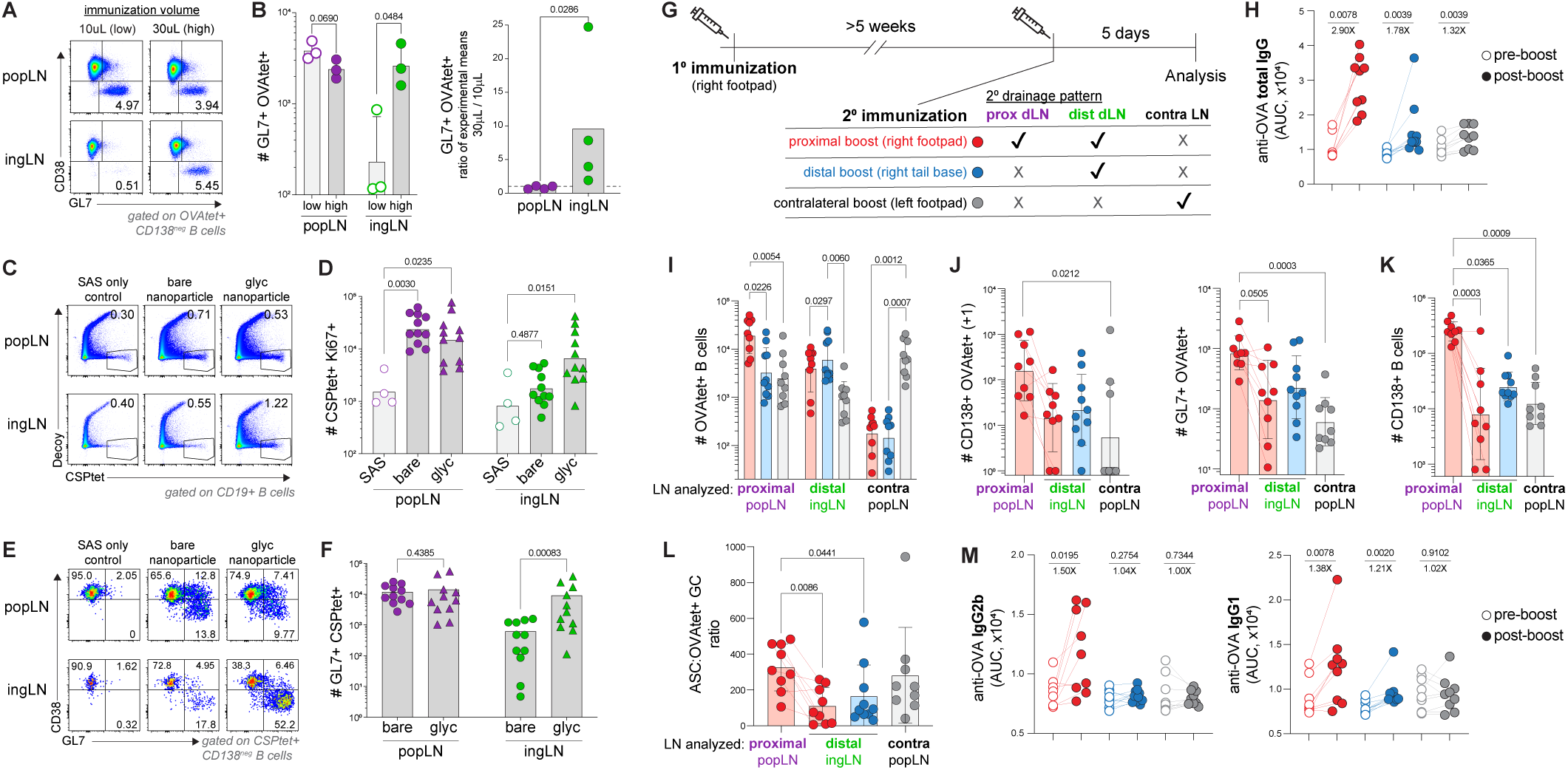
Vaccine design tunes B cell responses across the lymphatic network. A-B) B6 mice were footpad immunized with alum-absorbed OVA (10 μg) and LPS (10 μg) in either 10 μL or 30 μL volume; LNs analyzed day 9. A) Representative plots of OVA-tetramer+ CD19+ CD138^neg^B cells. B) Total number of OVA-tetramer+ GC B cells (left). Ratio of experimental means for total OVA-tetramer+ GC B cells after 30 μL versus 10 μL immunization. C-F) B6 mice immunized in the footpad or i.m. with Sigma Adjuvant System (SAS) plus non-glycosylated (bare) or glycosylated (glyc) nanoparticles displaying CSP antigen; LNs analyzed day 9. C, E) Representative plots of CSP-tetramer+ B cells. D, F) Flow quantification of Ki67+ activated (D) and GL7+ GC (F) CSP-tetramer+ B cells. G) Experimental setup for H-M. B6 mice were immunized in right footpad with SAS plus OVA (1° immunization). ≥5 weeks later mice were boosted (2° immunization) in right footpad (red), right tail base (blue), or left footpad (grey). LNs analyzed 5 days post-boost. Serum collected days −1 and 5 of boost. H) OVA-specific pan-IgG serum titers, pre-and post-boost. I-L) Flow quantification of total OVA-tetramer+ B cells (I), OVA-tetramer+ ASCs and GC B cells (J), total ASCs (K), and ratio of total ASCs to OVA-tetramer+ GC B cells (L). M) OVA-specific IgG2b (left) and OVA-specific IgG1 (right) serum titers pre-and post-boost. Two-tailed Mann-Whitney unpaired t-test (B, F), one-way ANOVA with Dunn’s multiple comparisons test (D, I-L), or Wilcoxon paired t-test (H, M). Connected points indicate LNs (J-L) or sera (H, M) from the same mouse, error bars represent s.d. Data pooled from ≥3 experiments.

We next asked whether intrinsic biochemical properties of the vaccine itself could similarly modulate B cell responses across the lymphatic network. Protein nanoparticle vaccines have emerged as powerful platforms to improve immunogenicity, and high-mannose N-linked glycosylation of nanoparticle immunogens has been shown to enhance localization and retention within FDC networks in dLNs,^47,48^ thereby improving humoral immunity.^49^ We therefore hypothesized that enhancing nanoparticle retention might preferentially augment responses within distal dLNs, where antigen availability is limiting. To test this, we compared LN responses elicited by non-glycosylated (bare) versus glycosylated nanoparticles displaying the *Plasmodium falciparum* circumsporozoite protein (CSP),^49^ the major surface antigen of the malaria parasite. While both nanoparticle formulations elicited robust CSP-specific B cell responses in proximal popLNs, glycosylation selectively enhanced responses within distal ingLNs (**Figures 5C**, **5D, and S6B**). This enhancement was driven primarily by expansion of the GC B cell compartment, with more modest effects on ASC numbers (**Figures 5E**, **5F, S6C, and S6D**), consistent with increased antigen localization within FDC networks.^47,48^

Together, these findings demonstrate that spatial organization of humoral immunity is programmable. Vaccine design features, including site-specific targeting of inflammation, injection volume, and nanoparticle engineering can be used to selectively tune the spatial distribution of B cell responses along the lymphatic chain.

### Lymphatic targeting during prime-boost immunization shapes recall responses

Prime-boost vaccination is widely used to enhance humoral immunity,^50^ and the location of booster immunization (ipsilateral versus contralateral) can strongly influence recall responses, partly due to the retention of memory B cells within dLNs.^51–53^ Whether multiple LNs within the same draining lymphatic network similarly harbor recall potential, however, has not been explored.

To address this, mice were primed in the right footpad and boosted >5 weeks later at one of three sites: the right footpad (ipsilateral proximal boost), the right tail base (ipsilateral distal boost), or the left footpad (contralateral boost) (**Figure 5G**). This strategy allowed us to assess whether early day 5 recall responses were shaped by the anatomical site of boost relative to primary immunization.

We found that boost location profoundly influenced the early recall responses in a hierarchical fashion. Ipsilateral proximal boosting produced the largest increase in antigen-specific serum titers, followed by ipsilateral distal boost, whereas contralateral boost elicited only a modest increase (**Figure 5H**). To investigate the basis of these differences, we assessed antigen-specific responses within individual LNs. Consistent with previous reports,^51–53^ ipsilateral proximal boosting elicited the strongest antigen-specific B cell response within the draining popLN characterized by antigen-specific GC responses as well as total and tetramer-binding ASCs (**Figures 5I-5K, red symbols**). Notably, antigen-specific responses were also significantly enhanced within the ipsilateral distal ingLN (**Figures 5I and 5J, red symbols**), albeit to a lesser extent, demonstrating that distal dLNs can participate in recall responses following same-site boost. Together, these data indicate ipsilateral proximal boosting promotes recall responses across multiple dLNs within the lymphatic network, and this distributed recall collectively contributes to enhanced systemic antibody titers.

Furthermore, directly targeting the distal ingLN via tail base boosting selectively enhanced responses at distal ingLN without affecting the proximal popLNs (**Figures 5I and 5J, blue symbols**), demonstrating that recall responses can be spatially directed to individual dLNs through site-directed booster immunization. Notably, both proximal and distal ipsilateral boosts promoted recall responses within distal ingLNs that skewed towards GC B cells over ASC responses, resulting in a decreased ASC:GC ratio in distal ingLNs compared to recall responses in proximal popLNs after ipsilateral proximal boost (**Figure 5L**). Such results indicate distal dLNs are key sites to consider for engagement of secondary GCs. In contrast, antigen-specific responses in the contralateral popLN were only observed following contralateral boosting, which were numerically reduced compared to ipsilateral popLN boost responses and primarily driven by ASCs rather than GC B cells (**Figures 5I-5K, grey symbols**). Together, these findings suggest that both ipsilateral proximal and distal dLNs harbor resident memory B cell populations capable of supporting rapid secondary responses, although the proximal dLN remains the most potent site of recall, whereas contralateral LNs are limited in their early recall capacity.

Finally, we ascertained whether recall responses also retained qualitative features established during priming, particularly isotype profiles. While direct flow staining for isotype classification proved technically challenging for early recall responses, serum antibody analysis revealed distinctions according to boost location. Antigen-specific serum IgG2b titers were only elevated with ipsilateral proximal boosting (**Figure 5M**), consistent with antigen-specific IgG2b responses restricted to proximal dLNs during priming (**Figures 4A-4C**). In contrast, antigen-specific serum IgG1 titers increased with both ipsilateral proximal and distal boosting (**Figure 5M**), mirroring the broader distribution of IgG1 class-switched responses established during priming (**Figures 4A-4C**). While contralateral boosting led to a modest increase in antigen-specific total IgG serum titers (**Figure 5H**), we did not detect a significant increase in either IgG2b nor IgG1 isotypes (**Figure 5M**), suggesting limited contribution of circulating IgG2b+ and IgG1+ memory B cells to these early responses.

Together, these findings suggest that immunological memory is distributed across multiple LNs within the draining lymphatic network and shapes the magnitude, cellular composition, and qualitative features of recall responses. These results further highlight that the anatomical location of booster immunization can be strategically leveraged to selectively engage individual LNs to modulate distinct features of vaccine-induced humoral immunity.

## DISCUSSION

Our study reveals a previously unrecognized spatial organization of humoral immunity across the lymphatic network. Rather than functioning as equivalent sites of B cell activation, individual dLNs generate distinct yet complementary immune responses that collectively determine the magnitude, quality, and diversity of humoral immunity. These differences arise from gradients of antigen and inflammatory cues established across the lymphatic chain, resulting in divergent microenvironments that differentially regulate Tfh differentiation, GC selection, ASC accumulation, isotype class switching, and recall responses. While this study was focused on vaccine-driven responses, similar lymphatics-based gradients likely exist in infection settings, helping to diversify immunological responses for both immediate and long-term host defense.

Previous studies demonstrated that adjuvant formulations can alter lymphatic flow and enhance delivery of vaccine component to distal dLNs, resulting in improved antibody responses in mice and non-human primates.^18,22^ Here, we extend this concept by demonstrating that additional vaccine design strategies, including site-specific LN targeting, injection volume, and nanoparticle engineering, can tune B cell responses across the lymphatic network, providing a proof-of-concept framework in which spatially encoded immune heterogeneity can be harnessed through vaccine design. Notably, across the various perturbations we tested, responses within proximal dLNs remain relatively stable, whereas responses within distal dLNs were preferentially modulated. These findings suggest that individual dLNs operate under distinct constraints, with proximal dLNs rapidly reaching maximal functional capacity under standard immunization conditions while distal dLNs are limited with regards to antigen availability, inflammatory cues, and early Tfh support, making them particularly amenable to perturbations via vaccine engineering.

Despite reduced antigen availability and delayed Tfh differentiation, distal dLNs supported productive GC responses that arose independently from those in proximal dLNs. Critically, proximal and distal dLNs exhibited distinct clonal dynamics, demonstrating that a diverse and affinity-matured B cell repertoire can emerge through multiple evolutionary trajectories within the same immune response. Proximal dLNs initially recruited a broad repertoire comprising high-and low-affinity populations, with select clones undergoing oligoclonal expansion, likely representing clonal jackpotting.^9,54^ In contrast, distal dLNs imposed more stringent selection thresholds during GC initiation, leading to early enrichment of high-avidity B cells and reduced clonal diversity from the outset. Delayed Tfh differentiation likely contributes to this early selection bottleneck, as GC-Tfh densities converged after GC establishment and provision of additional cognate CD4 T cell help selectively enhanced GC responses in distal LNs. Whether delayed Tfh differentiation reflects differences in antigen presentation, local cytokine environments, or other mechanisms remains an important question for future investigation. Importantly, our observations suggest a functional division of labor across the lymphatic network in which proximal dLNs preserve repertoire breadth, potentially enhancing protection against pathogenic variants,^55,56^ whereas distal dLNs preferentially contribute to high-affinity populations, including IgM+ memory B cells.

Another major finding was that robust ASC responses were preferentially found in proximal dLNs and were associated with a remodeled medullary compartment characterized by extensive expansion of capillary networks. Supporting past observations,^39,42^ ASCs within the medulla remained largely sessile, suggesting these regions represent specialized microenvironments for local ASC accumulation. How the remodeled medulla supports ASC persistence remains to be determined. One possibility is that the expanded capillary beds provide metabolic support, such as oxygen and glucose, akin to vascular niches that enable long-term ASC maintenance in the bone marrow.^57,58^ Notably, ASCs can produce VEGF and other factors that may promote endothelial cell proliferation,^59^ raising the possibility of a feed-forward circuit in which ASCs reinforce the vascular niche that supports their accumulation. In addition, ASCs are increasingly recognized to shape inter-and intra-clonal competition through antibody-feedback mechanisms.^60–62^ Thus, enhanced ASC responses in proximal dLNs raise the intriguing possibility that antibody feedback also influences GC evolution across the lymphatic network.

Finally, we demonstrated that the spatial organization of humoral immunity across the lymphatic network was maintained during recall responses. Given the diversity of memory B cells in phenotype, function, and tissue distribution,^63^ these findings suggest that spatial gradients also contribute to functional heterogeneity among memory B cell populations distributed across the lymphatic network. While previous studies have highlighted the importance of ipsilateral versus contralateral boosting,^51–53^ our study identifies an additional spatial axis in which selective targeting of individual LNs within the same lymphatic chain shapes early recall responses, including ASC differentiation, serum antibody magnitude and isotype composition, and secondary GC responses.

Altogether, our findings reveal that humoral immunity is spatially encoded, establishing the lymphatic network as both a determinant and programmable lever for vaccine design. Beyond diversifying responses, the ability to engage a greater number of LNs during vaccination may enhance recruitment and activation of rare broadly neutralizing antibody precursor B cells, an important consideration for protective immunity against HIV^64,65^ or influenza infection.^66,67^ Given that humans possess extensive lymphatic networks in which multiple draining lymph nodes participate in vaccine responses,^68^ these principles may provide new opportunities to spatially program humoral immunity toward broader and more effective responses.

## RESOURCE AVAILABILITY

### Lead contact

Requests for further information and materials should be directed to and will be fulfilled by Michael Y. Gerner.

### Materials availability

All materials generated in this study will be made available upon request to the lead contact.

### Data and code availability

The processed sequencing data in Figures 2 and 4 and S3 are available in 10.5281/zenodo.21249308, code available at https://github.com/rawlings-lab/lymphnode-bcrs. All sequences available at SRA under BioProject PRJNA1467187.

## ACKNOWLEDGEMENTS AND FUNDING.

Thank you to members of the Gerner Lab for assisting with experiments and helpful discussion. This work was supported by NIH grants R01AI134713 (M.Y.G) and T32AR007108 (J.L.C), the University of Washington Institute for Translational Immunology Postdoctoral Fellowship (J.L.C.), the Children’s Guild Association Endowed Chair in Pediatric Immunology and the Hansen Investigator in Pediatric Innovation Endowment (to D.J.R.). and the Gates Foundation grant number INV-043758 (N.P.K). The University of Washington Cell Analysis Facility Shared Resource Lab is supported in by part by NIH award 1S10OD024979-01A1.

## AUTHOR CONTRIBUTIONS

Conceptualization: J.L.C and M.Y.G. Investigation and analysis: J.L.C. and C.D.T. Methodology and resources: J.L.C., C.D.T., C.E.M., M.D.L., N.P.K, M.P., D.J.R., and M.Y.G. Funding acquisition: M.Y.G. Supervision: M.Y.G. Writing, original draft: J.L.C. and M.Y.G. Writing, review and editing: J.L.C., C.D.T., C.E.M., M.D.L., N.P.K., M.P., D.J.R., M.Y.G.

## SUPPLEMENTAL INFORMATION

**Figure S1 (related to Figure 1).**
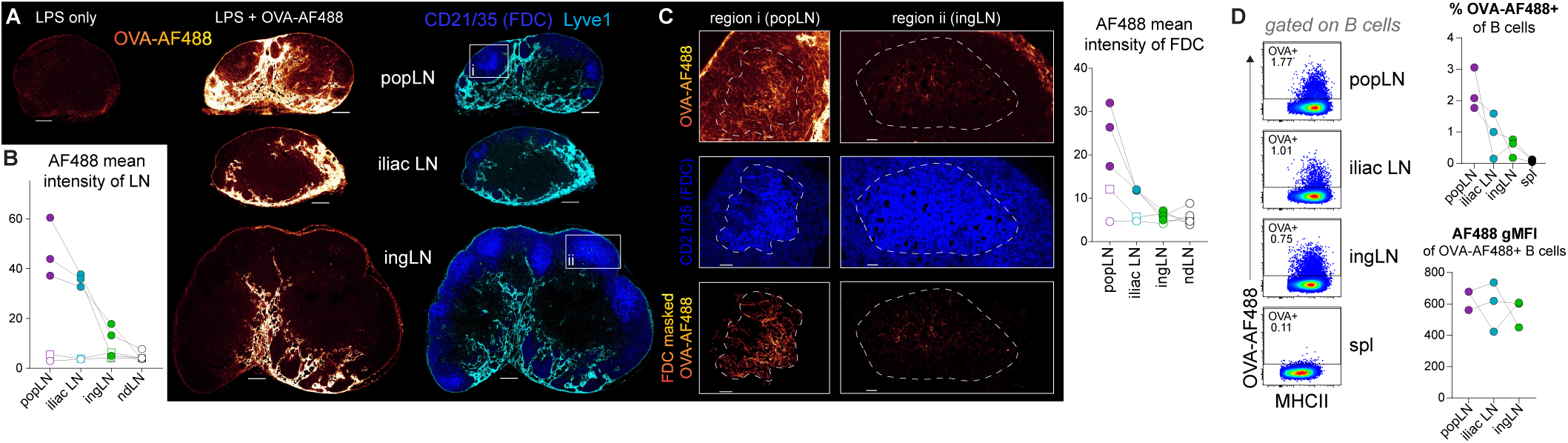
Antigen gradient across the draining lymphatic network established early after immunization. B6 mice were footpad immunized with OVA-AF488 plus LPS (A-C) or alum-absorbed OVA-AF488 and LPS (D); LNs analyzed 4 hours later. A) Representative confocal images of OVA-AF488 signal (look-up table (LUT)) with respect to CD21/35 and Lyve1 staining in draining LNs. Scale bar = 200 μm. B) Quantification of AF488 MFI per LN section. C) Representative zoom-in on FDC regions (left, middle). Quantification of AF488 MFI on FDC surfaces (right). Scale bar = 40 μm. D) Flow quantification of OVA-AF488+ B cells. Connected points indicate LNs from the same mouse. Open square = PBS control; open circle = LPS only (B, C).

**Figure S2 (related to Figure 1).**
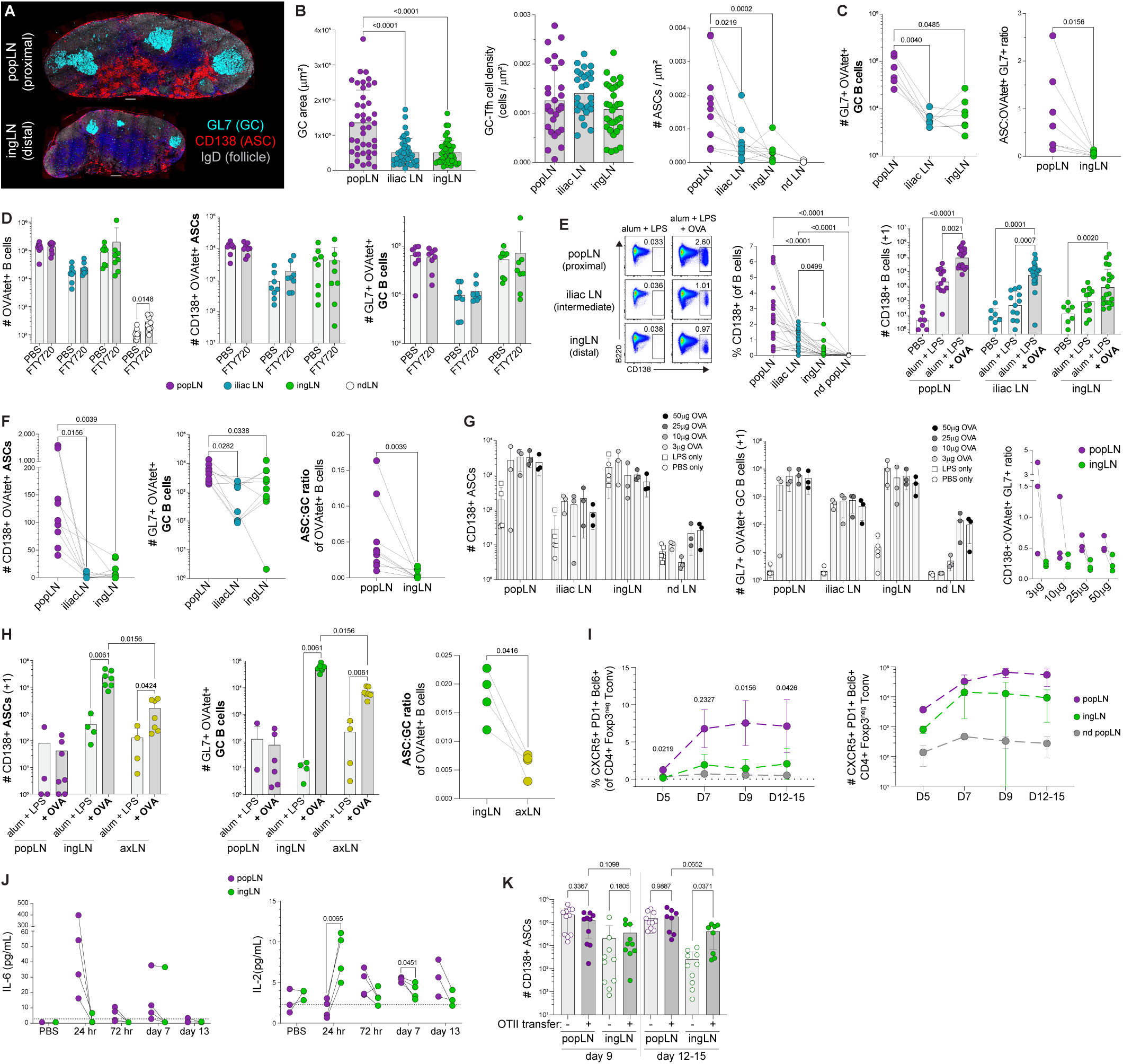
Antigen-specific B cell responses across the draining lymphatic network. A-E) B6 mice were footpad immunized with alum-absorbed OVA and LPS. A) Representative images of draining LNs 15 days post-immunization. B) Quantification of GC area (left), PD-1+ Tfh cell density within GC (middle), and IRF4+ CD138+ ASC density (right). C) Day 15 flow quantification of B cell responses. D) Day 9 quantification of B cell responses; +/− FTY720 administered on days 7-8. E) Flow quantification of ASCs; day 9. F-H) Day 9 quantification of responses following footpad immunization with Sigma Adjuvant System (SAS) plus OVA (F) or LPS plus OVA at indicated doses (G), or after tail base immunization with alum-absorbed OVA and LPS (H). I-K) Responses following footpad immunization with alum-absorbed OVA and LPS. I) Endogenous Tfh frequency (left) and number (right) at indicated timepoints. J) LEGENDplex cytokine screen of LN lysates at indicated timepoints post-immunization. Dotted line indicates limit of quantification. K) Total ASCs at indicated timepoints, +/− OT-II adoptive transfer. Two-tailed one-way ANOVA with Dunn’s (B, C left panel, E, I, K) or Tukey’s (F middle panel) multiple comparison, Wilcoxon paired-t-test (C right, F left and right, H left and middle panels, paired data), Mann-Whitney unpaired t-test (D, H left and middle), or paired t-test (H right, J). Connected points indicate LNs from same mice (B, C, E-H, J), error bars represent s.d. Data pooled from ≥2 experiments, except (J): displays three biological replicates from one experiment.

**Figure S3 (related to Figure 2).**
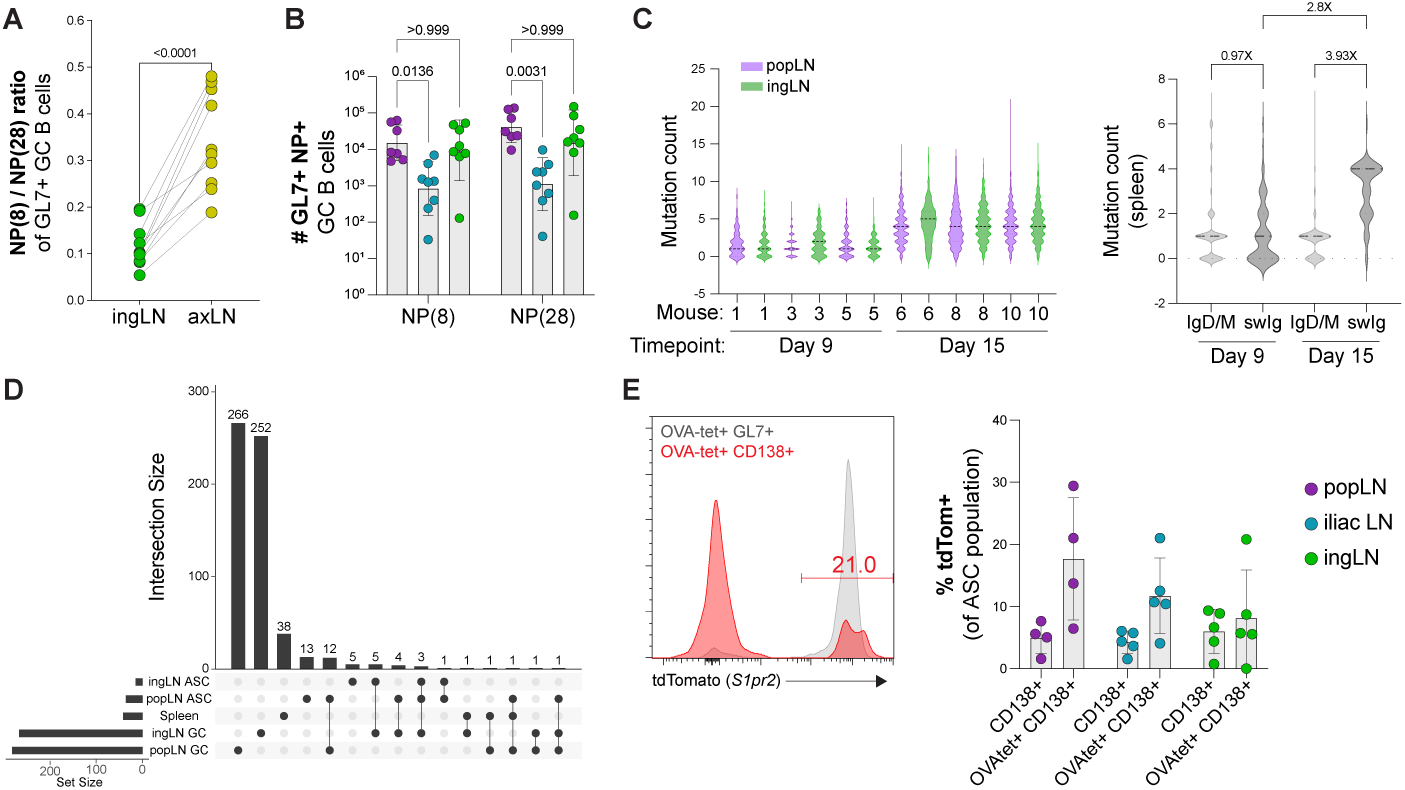
Antigen-specific repertoire analysis in proximal and distal draining LNs. A) Flow quantification of NP(8)+/NP(28)+ GC B cell ratio 9 days post tail base immunization with alum-absorbed NP-OVA and LPS. B) Flow quantification of total NP(8)-and NP(28)-binding GC B cells 15 days post footpad immunization with alum-absorbed NP-OVA and LPS. C-D) BCR repertoire analysis of sorted OVA-tetramer+ populations at days 9 and 15 post footpad immunization with alum-absorbed OVA and LPS. C) Total mutation count per sequence, by individual mouse and LN (left) or cumulative spleen data by isotype group (right). D) Representative UpSet plot displaying repertoires of indicated populations from a single mouse at day 15 post-immunization. Horizontal bars represent number of clones (set size) per population; vertical bars represent the set size per intersecting population denoted by black dots. E) Flow cytometry analysis of S1pr2-CreERt2 x Rosa-LSL-tdTom reporter mice following footpad immunization with alum-absorbed OVA and LPS. Tamoxifen treatment daily on days 6-8 post-immunization; LNs analyzed on day 9. Two-tailed paired t-test (A), or one-way ANOVA with Dunn’s (B, left panel) or Tukey’s (B, right panel) multiple comparisons test. Connected points indicate LNs from the same mouse (A, B), error bars represent s.d. Data are pooled from 2 experiments (A, B, E), or representative of or display three independent mice per group (C, D).

**Figure S4 (related to Figure 3).**
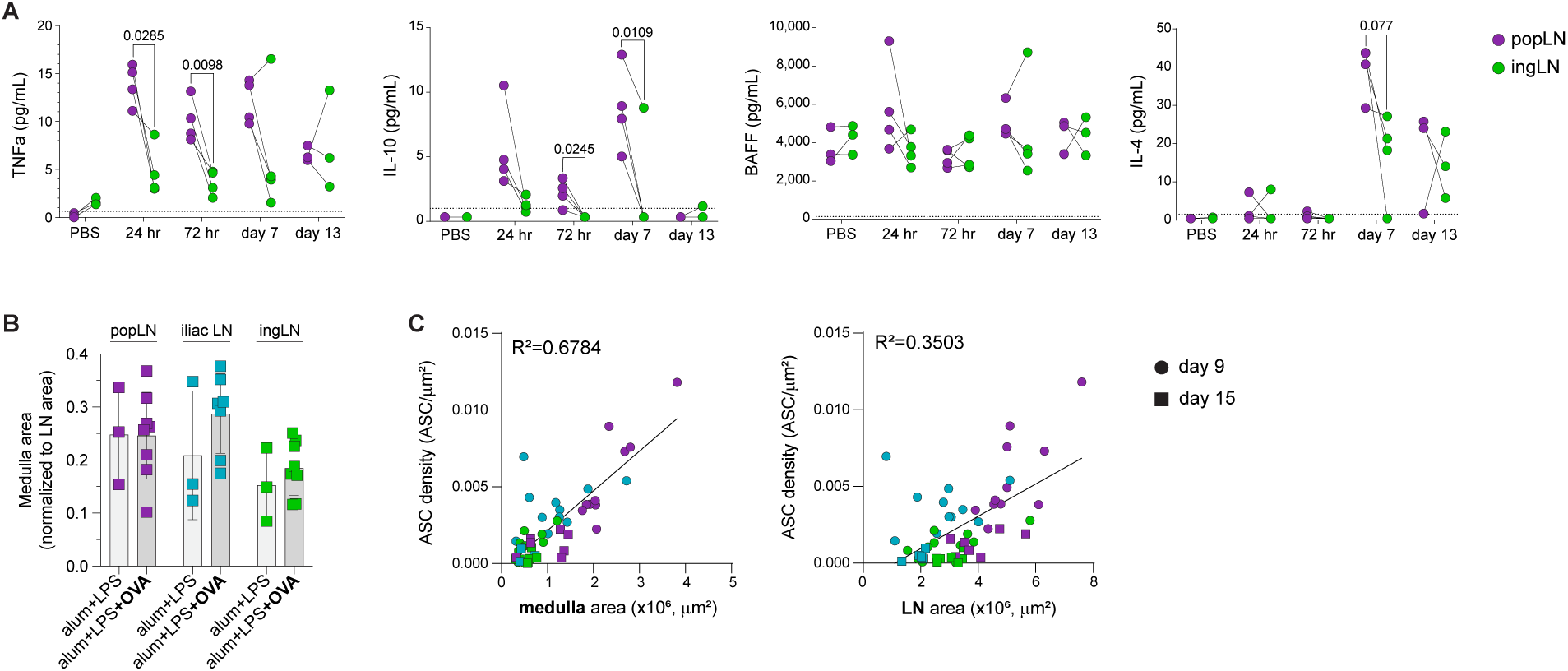
ASC responses are correlated with expanded LN medulla. B6 mice were footpad immunized with alum-absorbed OVA and LPS. A) LEGENDplex cytokine screen of LN lysates at indicated timepoints. Dotted line indicates limit of quantification. Connected dots indicate LNs from the same mouse. B-C) Imaging quantification. B) Medulla area normalized to total LN area; day 15. C) Correlation between IRF4+ CD138+ ASC density and medullary area (left) or total LN area (right) on day 9 (circles) or 15 (squares). Purple = popLN, aqua = iliac LN, green = ingLN. Linear regression performed and Goodness of Fit displayed as R^2^. Error bars represent s.d. Data are pooled from ≥2 experiments, except (A): three biological replicates from one experiment.

**Figure S5 (related to Figure 4).**
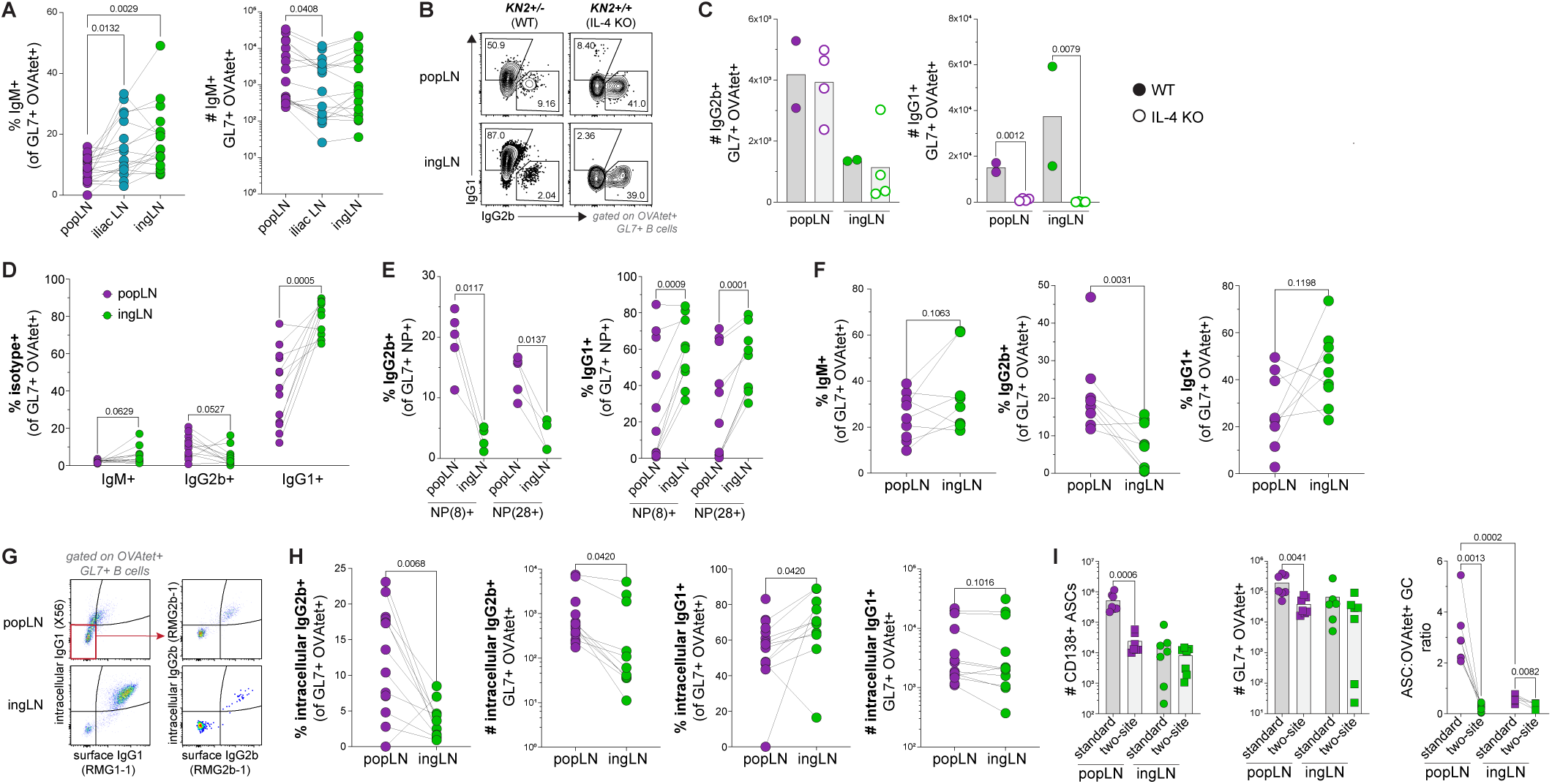
Distinct isotype profiles in proximal and distal draining LNs. A-E) Mice were footpad immunized with alum-absorbed OVA (A-D) or NP-OVA (E) and LPS. A) Flow analysis of B6 mice, 9 days post-immunization. B-C) KN2+/− WT or KN2+/+ IL-4 knock-out littermates were immunized; LNs analyzed day 9. B) Representative flow plots of OVA-tetramer+ GL7+ IgD^neg^ IgM^neg^ switched GC B cells. C) Quantification of OVA-tetramer+ GC B cells. D) GC B cell isotype analysis, B6 mice 12-15 days post-immunization. E) Isotype analysis of NP(8)+ and NP(28)+ GC B cells 9 days post-immunization. F) Isotype analysis 9 days post footpad immunization with SAS plus OVA. G-H) Flow analysis of intracellular and surface IgG2b and IgG1 staining, 9 days post footpad immunization with alum-absorbed OVA and LPS. G) Representative flow plots and H) quantification of OVA-tetramer+ GL7+ IgD^neg^ IgM^neg^ switched GC B cells. I) Mice were immunized according to experimental setup described in Figure 4D. Quantification of flow cytometry data 9 days post-immunization. Two-tailed one-way ANOVA with Tukey’s multiple comparisons test (A), Welch’s unpaired t-test (C, I right panel between immunization groups), paired t-test (D-F, I right panel between matched LNs of same mouse), Wilcoxon paired-test (H), or Mann-Whitney unpaired t-test (I left and middle panels). Connected points indicate LNs from the same mouse, error bars represent s.d. Data are pooled from ≥2 experiments, except (C) and IgG2b+ data in (E): one experiment.

**Figure S6 (related to Figure 5).**
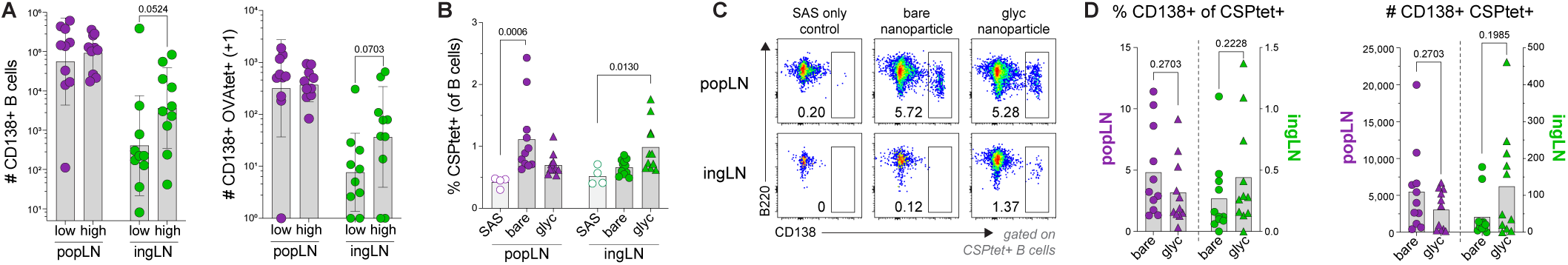
Vaccine design can tune B cell responses at distal draining LNs. A) Flow quantification of total CD138+ ASCs 9 days post footpad immunization with alum-absorbed OVA (10 μg) and LPS (10 μg) in either 10 μL or 30 μL volume. B-D) B6 mice were immunized in the footpad or i.m. with SAS plus non-glycosylated (bare) or glycosylated (glyc) nanoparticles displaying CSP antigen; LNs analyzed by flow cytometry 9 days later. B) Quantification of the frequency of CSP-tetramer+ cells of CD19+ B cells. C) Representative flow plots of total CD138+ ASCs. D) Quantification of total CD138+ ASCs in popLNs (purple, left y-axis) and ingLNs (green, right y-axis). Two-tailed Mann-Whitney unpaired t-test (A, D), or one-way ANOVA with Dunn’s multiple comparisons test (B). Error bars represent s.d. Data are pooled from ≥3 experiments.

## METHODS

### Mice

The following mice were purchased from The Jackson Laboratory: C57BL/6J mice (strain 000664), B6.Cg-*Gt(ROSA)26Sor^tm^*^14^*^(CAG-tdTomato)Hze^*/J (Ai14 tdTomato) mice (strain 007914), B6.Cg-Tg(Prdm1-EYFP)1Mnz/J (Blimp1-YFP) mice (strain 008828). CD45.1+ B6.Cg-Tg(TcraTcrb)425Cbn/J (OT-II) mice were obtained from a donating investigator (P.J. Fink, University of Washington, Seattle, WA). S1PR2-ERT2Cre^+^ mice were kindly provided by Tomohiro Kurosaki (RIKEN Center for Integrative Medical Sciences, Yokohama, Japan) and crossed to Ai14 tdTomato mice to generate S1PR2-fate reporter mice. KN2 (IL-4 KO) mice on the B6 background were provided by Dr. Markus Mohrs (Trudeau Institute). Mice were 6-16 weeks of age at time of immunization, except where noted. For immunizations, male wild-type mice, female Blimp1-YFP reporter mice, female and male S1pr2-fate reporter mice, and female and male KN2 mice were used. For adoptive T cell transfer experiments, 7-19 week old OT-II female and male mice were used. All mice were maintained in specific pathogen-free conditions at an Association for Assessment and Accreditation of Laboratory Animal Care-accredited facility at the University of Washington, South Lake Union campus. All procedures were approved by the University of Washington Institutional Animal Care and Use Committee.

### Immunizations

All immunizations were administered intradermally (i.d.) into the hind footpad and/or subcutaneously (s.c.) at the tail base or intramuscularly (i.m.) into the thigh in a volume of 20 μL, unless otherwise noted. For alum-based immunizations, 10 μg LPS from *E. coli* O111:B4 (Sigma-Aldrich) with or without 10 μg endotoxin-free Ovalbumin (OVA) (Invivogen), unless otherwise noted, were mixed in PBS and absorbed to aluminium hydroxide gel (alum) (Invivogen) at a 1:1 ratio. In some experiments, 10 μg NP-OVA (Biosearch Technologies, conjugation ratio of 20) was used as antigen. For antigen tracking experiments in Figure S1, 20 μg OVA-AF488 (Invitrogen) was used. For some experiments, Sigma Adjuvant System (SAS, Sigma-Aldrich) was mixed at a 1:1 ratio with 10 μg endotoxin-free OVA or 3 μg total nanoparticle protein (1.02 μg CSP antigen).

### *in vivo* treatments

For adoptive transfers, naïve CD45.1+ OT-II T cells were isolated from LNs and spleens using the naïve CD4+ T cell isolation kit (Miltenyi Biotec). 1-2×10^5^ naïve OT-II T cells were transferred into CD45.2+ C57BL/6J hosts intravenously one day prior to immunization. To block lymphocyte egress, mice were treated with sphingosine-1-phosphate receptor agonist FTY720 (Cayman Chemical) at a concentration of 1 μg per gram of mouse weight on days 7 and 8 post-immunization via intraperitoneal injection. To fate-map S1pr2-expressing cells, S1pr2-fate reporter mice were treated daily with 1 mg tamoxifen (Sigma-Aldrich) in corn oil (Sigma-Aldrich) on days 6-8 post immunization via intraperitoneal injection.

### Confocal and intravital two-photon microscopy

For confocal imaging, isolated LNs were fixed using BD Cytofix (BD Biosciences) diluted 1:3 with PBS for 15-24 hours at 4°C and then dehydrated with 30% sucrose for 15–48 hours at 4°C. Tissues were then embedded in OCT compound (Tissue-Tek) and stored at −80°C. LNs were sectioned on a Thermo Scientific Microm HN550 cryostat into 20 μm sections and were then stained with fluorescently conjugated antibodies at room temperature or 4°C for 6–18 hours. Slides were washed with 0.1 M Tris buffer and cover-slipped using Fluoromount G mounting media (SouthernBiotech). A Leica SP8 tiling confocal microscope equipped with a 40X 1.3 NA oil objective was used for confocal image acquisition. For intravital two-photon microscopy, mice were anesthetized using an isoflurane vaporizer and popliteal LNs were surgically exposed. Imaging was performed using a Leica SP8 microscope outfitted with a Chameleon laser and a 20X 1.0 NA water immersion objective. For vascular labeling, mice were intravenously administered 300 μg Dextran (70,000 molecular weight; Invitrogen; FITC conjugate) 1–6 hours prior to imaging. For experiments imaging Blimp1-YFP reporter mice, 1 μg of anti-FAS-PE antibody was injected i.d. in the footpad 4 hours prior to imaging to identify GCs and also used to visualize antibody-capturing phagocytes. To label ASCs in C57BL/6J mice, 1 μg CD138-PE antibody was injected intradermally to the footpad 3–7 hours prior to imaging. All acquired raw imaging data was processed and analyzed in Imaris (Bitplane).

### Microscopy analysis

Multiparameter confocal images were corrected for fluorophore spillover using the Leica Channel Dye Separation module. All images were visualized and analyzed in Imaris (Bitplane). ASC objects were created on IRF4+ CD138+ cells using the surface object creation wizard in Imaris. Total LN area was manually defined using all channels to delineate the LN boundary and represented as a surface. LN medulla regions were manually identified as the region outside of T cell zones (identified by CD3 or CD4 staining and associated with Collagen IV+ FRC conduits) and B cell follicles (identified by B220 or IgD staining), enriched for Lyve1+ lymphatics and Collagen IV+ stromal structures within the LN hilus, and represented as a surface. GC regions were manually identified using combined staining for Ki67+ Bcl6+ and/or GL7+ regions in B cell follicles and represented as a surface. GC-Tfh objects were created on PD-1+ signal within GC regions, and manually confirmed as CD4+ Bcl6+. For data presented in Figure S1C, FDC objects were generated on CD21/35^HIGH^ B220^LOW/NEG^ staining using the surface object creation wizard, and AF488 signal was masked outside of FDC objects. For intravital two-photon microscopy, ASC objects were created on Blimp1-YFP+ FAS^NEG^ non-phagocytic cells or CD138-PE+ cells using the spot object creation wizard.

### Tetramer generation

Recombinant full-length PfCSP (SAmut-CSP)^69^ was expressed using HEK293F cells (Thermo Scientific) and then column purified using the Ni-NTA kit (Millipore Sigma). Endotoxin-free OVA (Invivogen) and purified recombinant SAmut-CSP proteins were biotinylated using the BirA500 protein ligase reaction kit (Avidity), according to the manufacturer’s instructions. Biotinylated proteins were then tetramerized as previously described.^70^ Briefly, biotinylated Ovalbumin and PfCSP proteins were incubated with streptavidin-APC or streptavidin-PE (Agilent), respectively, at room temperature for 30 minutes. The tetramer fraction was centrifuged in a 100 kDa Amicon filter (Millipore) to filter out non-tetramerized protein. A decoy reagent to exclude nonspecific binding to the OVA-APC tetramer was made by conjugating SA-APC to DyLight 755 using a DyLight 755 antibody labeling kit (Thermo Scientific), washing and removing any unbound DyLight 755, and incubating with an excess of an irrelevant biotinylated His-tagged class II–associated invariant chain peptide. A decoy reagent to exclude nonspecific binding to the PfCSP-PE tetramer was made by conjugating SA-PE to AF647 using an AF647 protein labeling kit (Thermo Scientific), washing and removing any unbound AF647, and incubating with an excess of an irrelevant biotinylated His-tagged class II–associated invariant chain peptide.

### Cell isolation and flow cytometry

LNs were mechanically disrupted and filtered through Nitex mesh (Genesee Scientific). For B cell tetramer staining, single cell suspensions were stained with decoy tetramer at a concentration of 10 nM in staining buffer (PBS containing 2% FBS and Fc block (2.4G2)) for 25 minutes at room temperature. OVA-APC or CSP-PE tetramer was added at a concentration of 10 nM, along with surface antibodies and incubated for 30 minutes on ice. For CXCR5 staining, cells were incubated with CXCR5-biotin for 45 minutes at room temperature. For intracellular staining using the FOXP3 Fix/Perm kit (Invitrogen), cells were fixed for 30 minutes on ice followed by incubation with intracellular antibodies for 30 minutes on ice, except for Foxp3 which was stained overnight at 4°C. For OVA-specific B cell enrichment of spleen samples, single-cell suspensions of RBC-lysed splenocytes were stained with decoy tetramer, as described above. OVA-APC tetramer was then added at a concentration of 10 nM and incubated for 30 minutes on ice. Cells were washed, incubated with anti-APC magnetic beads for 30 minutes on ice, and enriched for using magnetized LS columns (Miltenyi Biotec). All bound cells were stained with surface antibodies, followed by fixation and intracellular antibody staining, as described above. To determine the population avidity of NP-specific GC B cells, single cell suspensions of individual LNs were divided into two equal fractions, each incubated with surface antibodies including either NP(8) or NP(28) staining probes (Biosearch Technologies), followed by fixation and intracellular antibody staining, as described above. Flow cytometry data was acquired through the University of Washington, Cell Analysis Facility Shared Resource Lab, using the Cytek Aurora or BD Symphony A3 cytometer. Data was analyzed using FlowJo software. Cell sorting was performed using the BD Aria 3 Cell Sorter.

### Lymph node cytokine analysis

To quantify cytokine levels, draining LNs were lysed at indicated timepoints post-immunization using a Precellys CK14 lysing kit (Berin Corp) in 1X Pierce protease inhibitor (Thermo Scientific), and concentrated using a 10 kDa Amicon filter (Millipore). Samples were then processed with a LEGENDPlex 13-plex mouse B cell panel (Biolegend) according to manufacturer’s instructions and analyzed by flow cytometry on a BD CantoRUO (BD Biosciences). The data were analyzed using LEGENDplex software (BioLegend).

### anti-OVA serum ELISA

High binding 96-well plates (Corning) were coated with 50 μL per well of 5 μg/mL OVA diluted in PBS and incubated overnight at 4°C. Plates were washed with PBS containing 0.05% Tween-20 (PBS-T), then blocked with blocking buffer (PBS-T and 3% w/v non-fat milk powder) for two hours at room temperature. After washing with PBS-T, sera were diluted in blocking buffer and 50 μL per well was added and incubated for 1-2 hours at room temperature. After washing with PBS-T, bound antibodies were detected with 1:1000 dilution of biotinylated anti-mouse polyclonal IgG (BioLegend), monoclonal IgG2b (BioLegend), or monoclonal IgG1 (BioLegend) diluted in blocking buffer. After 1 hour at room temperature, plates were washed and then incubated with SA-HRP diluted in blocking buffer at 1:1000. After 30 minutes at room temperature, plates were washed five times with PBS-T, and developed using 50 μL 1xTMB (Thermo Scientific) and reactions quenched with 100 μL of 1M HCL. Absorbance at 450 nm was measured using an Epoch Microplate reader (Biotek) and values plotted and Area Under Curve (AUC) calculated using GraphPad Prism Software.

### CSP nanoparticle assembly and generation

CSP nanoparticle immunogens RT.2-I53-50 and RT.2-I53-50-3gly were assembled and characterized as previously described.^49^ Briefly, purified RT.2-I53-50A or RT.2-I53-50A-3gly trimer (non-glycosylated or glycosylated, respectively) was mixed with I53-50B.4PT1 pentamer at a 1.1:1 molar ratio^71^ in assembly buffer (25 mM Tris, 150 mM NaCl, 5% glycerol, pH 8.0) at 20–50 µM total protein and incubated overnight at 4°C with rocking. Assemblies were purified from residual free component by SEC on a Superose 6 Increase 10/300 GL column (Cytiva) in assembly buffer, pooling fractions at the expected elution volume for fully formed particles. Purity was assessed by SDS-PAGE and concentration by UV-vis (Agilent Cary 3500). Nanoparticle assembly was confirmed by negative-stain electron microscopy. For glycan analysis, N-linked glycosylation of the glycosylated trimer and corresponding nanoparticle was confirmed by glycosidase gel shift (PNGase F and Endo H, NEB) and site-specific glycan profiles at each sequon were determined by LC-MS/MS on an Orbitrap Ascend Tribrid (Thermo Fisher) to confirm the presence of high-mannose glycans, as previously described.^49^

### BCR-sequencing and repertoire analysis

#### Library Generation

cDNA was generated from bulk sorted cells using a Template Switching RT Enzyme Mix (NEB) and the manufacturer’s protocol for Smart-seq at double volume per reaction. To account for larger cell numbers greater than 1000, a longer 17 base barcoded Template Switch Oligo (TSO) was used, and a shorter 8 base barcoded TSO was used for counts less than 1000. Reactions used a poly-T primer that shared the sequence of the TSOs and allowed for subsequent smart-seq amplification of all cDNA. A 1/3 portion of the barcoded cDNA was used in a qPCR reaction to determine the optimal number of cycles before plateau for each reaction. Remaining barcoded cDNA was then amplified accordingly. To amplify BCRs, a multiplex PCR was performed on each bulk sample using pooled constant region primers for IgM, IgG, IgA, IgD, IgK, and IgL and sample-barcoded forward primers for the universal template switch region. For PacBio Revio sequencing, samples were pooled and library prepped according to manufacturer’s protocols. For Illumina sequencing, samples were pooled and library prepped using enzymatic fragmentation and ligation with an anchored multiplex PCR approach^72^ to retain sample and transcript barcodes. Indexed library preps were run on an Illumina NovaSeq X at 300 cycles.

#### Sequence Processing

Individual samples were demultiplexed by sample-barcoded sequences. For Illumina sequences, UMI-tools^73^ was used to create whitelists of transcript barcodes for each sample after manually determining the barcode counts from the knee in the rank abundance plots. TRUST4^74^ was run for each sample along with the barcode whitelist to create BCR contigs. Contigs were submitted to IMGT/HighV-Quest (GENE-DB Version 3.1.42) for annotation and nucleotide mutation counts were parsed using an in-house python script from the output tables.

Productive sequences for each sample were collapsed by V-gene, J-gene, nucleotide junction and amino acid mutations. For PacBio Revio long-read data, pRESTO (version 0.7.9)^75^ was used to mark UMIs that were filtered against a whitelist. Concatemers were removed with a custom Python script using Biopython (version 1.87).^76^ Sequences within identical 17 bp UMI groups were aligned using MUSCLE (version 3.8),^77^ and molecular consensus sequences were generated by collapsing groups with a read support of n >= 2. All sequences were aligned to reference IMGT databases using IgBLAST (version 1.6).^78^ Some barcode collisions were observed in the 8 bp UMIs and mitigated in R using tidyverse tools (version 1.2.1)^79^ by clustering CDR3 sequences with an indel distance of 1 using the stringdist package^80^ and removing sequences with a read count < 20 and < 6% of the maximum cluster count. These sequences were grouped by junction, C-gene, and sample; junctions accounting for < 1.5% of the maximum read count for that specific junction across cross-samples were removed. Sequences with a junction length not divisible by 3 were discarded. Leader sequences were trimmed, productive full-length sequences were collapsed to unique VDJC, and sequences containing J-region indels identified by CIGAR string were removed. Processed PacBio datasets were then integrated with the Illumina dataset for unified analysis.

#### Sequence Analysis

To quantify total mutation burden, sequences were length-filtered to ensure complete coverage of all complementarity-determining regions (CDRs), and mismatches and insertion/deletion (indel) events were summed by parsing V-and J-segment CIGAR strings. Inter-repertoire overlap of unique nucleotide junctions was evaluated per individual mouse across tissue organs and cell populations, with intersections visualized via the UpSetR package (version 1.4.0).^81^ Isotype distributions were assessed by tabulating sequence counts stratified by day, organ, cell population, and constant (C) gene call. Repertoire diversity metrics were computed using the immunarch package (version 0.10.3)^82^ compiled as bulk AIRR-seq data. Repertoire richness was estimated using the airr_diversity_chao1 function, and clonal dominance was calculated as D50 indices via the airr_diversity_dxx function with the percentage parameter set to 50.

#### Clonotype Assignment and Network Diagrams

Nearest-neighbor distances were calculated across the nucleotide junction region using a length-normalized Hamming distance model grouped by V and J gene calls (shazam 1.3.2).^83^ Based on the distance distribution histogram, a threshold of 0.04 was selected. Hierarchical clonal clustering was executed (scoper1.4.0)^33^ using the nucleotide sequence method with length normalization under the 0.04 threshold, partitioned strictly by individual mouse. For germline reconstruction, mouse V, D, and J germline reference databases were loaded from IMGT using dowser (version 2.4.1).^84^ Clonal germline sequences were generated for each assigned clone based on the sequence_alignment field and reference alignments while maintaining independent mouse stratification. Individual clonal trees were generated using buildPhylipLineage (via PHYLIP dnapars in alakazam 1.4.3)^83^ or initialized as single-vertex graphs for singletons. Clonotype graphs were merged into sample-specific networks across mouse and organ combinations using an attribute-preserving igraph::union routine. Networks were visualized using igraph 2.3.1 (R package version 2.3.1, https://CRAN.R-project.org/package=igraph).^85^ Vertices with multi-isotype profiles due to sequence collapsing were rendered as pie charts proportional to isotype frequencies. Single-isotype nodes were assigned shapes based on cell population (GC as circles; ASC as squares), with circle morphology prioritized in multi-population collapsed singletons. Node fill colors mapped directly to constant (C) gene classifications, with inferred nodes rendered in white.

### Statistical analysis

All statistical analyses were conducted using GraphPad Prism Software. Data points represent independent samples.

